# A Bayesian Mixture Modelling Approach For Spatial Proteomics

**DOI:** 10.1101/282269

**Authors:** Oliver M. Crook, Claire M. Mulvev, Paul D.W. Kirk, Kathryn S. Lillev, Laurent Gattot

## Abstract

**Abstract:** Analysis of the spatial sub-cellular distribution of proteins is of vital importance to fully understand context specific protein function. Some proteins can be found with a single location within a cell, but up to half of proteins may reside in multiple locations, can dynamically re-localise, or reside within an unknown functional compartment. These considerations lead to uncertainty in associating a protein to a single location. Currently, mass spectrometry (MS) based spatial proteomics relies on supervised machine learning algorithms to assign proteins to sub-cellular locations based on common gradient profiles. However, such methods fail to quantify uncertainty associated with sub-cellular class assignment. Here we reformulate the framework on which we perform statistical analysis. We propose a Bayesian generative classifier based on Gaussian mixture models to assign proteins probabilistically to sub-cellular niches, thus proteins have a probability distribution over sub-cellular locations, with Bayesian computation performed using the expectation-maximisation (EM) algorithm, as well as Markov-chain Monte-Carlo (MCMC). Our methodology allows proteome-wide uncertainty quantification, thus adding a further layer to the analysis of spatial proteomics. Our framework is flexible, allowing many different systems to be analysed and reveals new modelling opportunities for spatial proteomics. We find our methods perform competitively with current state-of-the art machine learning methods, whilst simultaneously providing more information. We highlight several examples where classification based on the support vector machine is unable to make any conclusions, while uncertainty quantification using our approach provides biologically intriguing results. To our knowledge this is the first Bayesian model of MS-based spatial proteomics data.

**Author summary:** Sub-cellular localisation of proteins provides insights into sub-cellular biological processes. For a protein to carry out its intended function it must be localised to the correct sub-cellular environment, whether that be organelles, vesicles or any sub-cellular niche. Correct sub-cellular localisation ensures the biochemical conditions for the protein to carry out its molecular function are met, as well as being near its intended interaction partners. Therefore, mis-localisation of proteins alters cell biochemistry and can disrupt, for example, signalling pathways or inhibit the trafficking of material around the cell. The sub-cellular distribution of proteins is complicated by proteins that can reside in multiple micro-environments, or those that move dynamically within the cell. Methods that predict protein sub-cellular localisation often fail to quantify the uncertainty that arises from the complex and dynamic nature of the sub-cellular environment. Here we present a Bayesian methodology to analyse protein sub-cellular localisation. We explicitly model our data and use Bayesian inference to quantify uncertainty in our predictions. We find our method is competitive with state-of-the-art machine learning methods and additionally provides uncertainty quantification. We show that, with this additional information, we can make deeper insights into the fundamental biochemistry of the cell.

## 1 Introduction

Spatial proteomics is an interdisciplinary field studying the localisation of proteins on a large-scale, Where a protein is localised in a cell is a fundamental question, since a protein must be localised to its required sub-cellular compartment to interact with its binding partners (for example, proteins, nucleic acids, metabolic substrates) and carry out its function (Gibson, 2009), Furthermore, mis-localisations of proteins are also critical to our understanding of biology, as aberrant protein localisation have been implicated in many pathologies (Olkkonen and Ikonen, 2006; Luheshi *et al.*, 2008; Laurila and Vihinen, 2009; De Matteis and Luini, 2011; Cody *et al.*, 2013), including cancer (Kau *et al.*, 2004; Rodriguez *et al.*, 2004; Latorre *et al.*, 2005; Shin *et al.*, 2013) and obesity (Siljee *et al.*, 2018).

Sub-cellular localisations of proteins can be studied by high-throughput mass spectrometry (MS) (Gatto *et al.*, 2010), MS-based spatial proteomics experiments enable us to confidently determine the sub-cellular localisation of thousands of proteins within in a cell (Christoforou *et al.*, 2016), given the availability of rigorous data analysis and interpretation (Gatto *et al.*, 2010).

In a typical MS-based spatial proteomics experiment, cells first undergo lysis in a fashion which maintains the integrity of their organelles. The cell content is then separated using a variety of methods, such as density separation (Dunkley *et al.*, 2006; Christoforou *et al.*, 2016), differential centrifugation (Itzhak *et al.*, 2016), free-flow electrophoresis (Parsons *et al.*, 2014), or affinity purification (Heard *et al.*, 2015), In LOPIT (Dunkley *et al.*, 2004, 2006; Sadowski *et al.*, 2006) and *hyper*LOPIT (Christoforou *et al.*, 2016; Mulvey *et al.*, 2017), cell lysis is proceeded by separation of the content along a density gradient. Organelles and macro-molecular complexes are thus characterised by density-specific profiles along the gradient (De Duve and Beaufay, 1981), Discrete fractions along the continuous density gradient are then collected, and quantitative protein profiles that match the organelle profiles along the gradient, are measured using high accuracy mass spectrometry (Mulvey *et al.*, 2017).

The data are first visualised using principal component analysis (PCA) and known sub-cellular compartments are annotated (Breckels *et al.*, 2016a), Supervised machine learning algorithms are then typically employed to create classifiers that associate un-annotated proteins to specific organelles (Gatto *et al.*, 2014a), as well as semi-supervised methods that detect novel sub-cellular clusters using both labelled and un-labelled features (Breckels *et al.*, 2013), More recently, a state-of-the-art transfer learning (TL) algorithm has been shown to improve the quantity and reliability of sub-cellular protein assignments (Breckels *et al.*, 2016b), Applications of such methods have led to organelle-specific localisation information of proteins in plants (Dunkley *et al.*, 2006), *Drosophila* (Tan *et al.*, 2009), chicken (Hall *et al.*, 2009), human cell lines (Breckels *et al.*, 2013), mouse pluripotent embryonic stem cells (Christoforou *et al.*, 2016) and cancer cell lines (Thul *et al.*, 2017).

Classification methods which have previously been used include partial least squares discriminate analysis (Dunkley *et al.*, 2006), K nearest neighbours (Groen *et al.*, 2014), random forests (Ohta *et al.*, 2010), naive Bayes (Xikolovski *et al.*, 2012), neural networks (Tardif *et al.*, 2012) and the support vector machine amongst others (see Gatto *et al.* (2014a) for an overview). Although these methods have proved successful within the field they have limitations. Typically, such classifiers output an assignment of proteins to discrete preannotated sub-cellular locations. However, it is important to note that half the proteome cannot be robustly assigned to a single sub-cellular location, which may be a manifestation of proteins in so far uncharaterised organelles or proteins that are distributed amongst multiple locations. These factors lead to uncertainty in the assignment of proteins to sub-cellular localisations, and thus quantifying this uncertainty is of vital importance (Kirk *et al.*, 2015).

To overcome the task of uncertainty quantification, this article presents a probabilistic generative model for MS-based spatial proteomics data. Our model posits that each annotated sub-cellular niche can be modelled by a multivariate Gaussian distribution. Thus, the full complement of annotated proteins is captured by a mixture of multivariate Gaussian distributions. With the prior knowledge that many proteins are not captured by known sub-cellular niches, we augment our model with an outlier component. Outliers are often dispersed and thus this additional component is described by a heavy-tailed distribution: the multivariate Student’s t-distribution, leading us to a T Augmented Gaussian Mixture model (TAGM).

Given our model and proteins with known location, we can probabilistically infer the sub-cellular localisation of thousands of proteins. We can perform inference in our model by finding *maximum a posteriori* (MAP) estimates of the parameters. This approach returns the probability of each protein belonging to each annotated sub-cellular niche. These posterior localisation probabilities can then be the basis for classification. In a more sophisticated, fully Bayesian approach to uncertainty quantification, we can additionally infer the entire posterior distribution of localisation probabilities. This allows the uncertainty in the parameters in our model to be reflected in the posterior localisation probabilities. We perform this inference using Markov-chain Monte-Carlo methods; in particular, we provide an efficient collapsed Gibbs sampler to perform inference.

We perform a comprehensive comparison to state-of-the-art classifiers to demonstrate that our method is reliable across 19 different spatial proteomics datasets and find that all classifiers we considered perform competitively. To demonstrate the additional biological advantages our method can provide, we apply our method to a *hyper*LOPIT dataset on mouse pluripotent embryonic stem cells (Christoforou *et al.*, 2016), We consider several examples of proteins that were unable to be assigned using traditional machine-learning classifiers and show that, by considering the full posterior distribution of localisation probabilities, we can draw meaningful biological results and make powerful conclusions. We then turn our hand to a more global perspective, visualising uncertainty quantification for over 5,000 proteins, simultaneously. This approach reveals global patterns of protein organisation and their distribution across sub-cellular compartments.

We make extensive use of the R programming language (R Core Team, 2017) and existing MS and proteomics packages (Gatto and Lilley, 2012; Gatto *et al.*, 2014b), We are highly committed to creating open software tools for high quality processing, visualisation, and analysis of spatial proteomics data. We build upon an already extensive set of open software tools (Gatto *et al.*, 2014b) as part of the Bioconductor project (Gentleman *et al.*, 2004; Huber *et al.*, 2015) and our methods are made available as part of this project.

## 2 Results

### 2.1 Application to mouse pluripotent embryonic stem cell data

We model mouse pluripotent embryonic stem cell (E14TG2a) data (Christoforou *et al.*,), which contains quantitation data for 5032 proteins. This high-resolution map was produced using the *hyper*LOPIT workflow (Mulvey *et al.*, 2017), which uses a sophisticated sub-cellular fractionation scheme. This fractionation scheme is made possible by the use of Tandem Mass Tag (TMT) 10-plex and high accuracy TMT quantification was facilitated by using synchronous precursor selection MS3 (SPS-MS3) (McAlister *et al.*, 2014), which reduces well documented issues with ratio distortion in isobaric multiplexed quantitative proteomics (Ting *et al.*, 2011), The data resolves 14 sub-cellular niches with an additional chromatin preparation resolving the nuclear chromatin and non-chromatin components. Two 10 gradient. We defined gold standard organelle markers as those with unambiguous single annotation (Gatto *et al.*, 2014a), A protein marker list for the mouse pluripotent embryonic stem cells was manually curated using information from the UniProt database, the Gene Ontology and the literature, as was performed in Christoforou *et al.* (2016), The following section applies our statistical methodology to these data and we explore the results.

#### 2.1.1 Maximum a posteriori prediction of protein localisation

This section applies the TAGM model to the mouse pluripotent embryonic stem cell data, by deriving MAP estimates for the model parameters and using these for prediction. Visualisation is important for data analysis and exploration, A simple way to visualise our model is to project probability ellipses onto a PCA plot. Each ellipse contains a proportion of total probability of a particular multivariate Gaussian density. The outer ellipse contains 99% of the total probability whilst the middle and inner ellipses contain 95% and 90% of the probability respectively Visualising only the first two principal components can be misleading, since proteins can be more (or less) separated in subsequent principal components. We visualise the first two principal components along with the first and fourth principal components as a representative example. For the TAGM model, we derive probability ellipses from the MAP estimates of the parameters.

**Figure 1:**
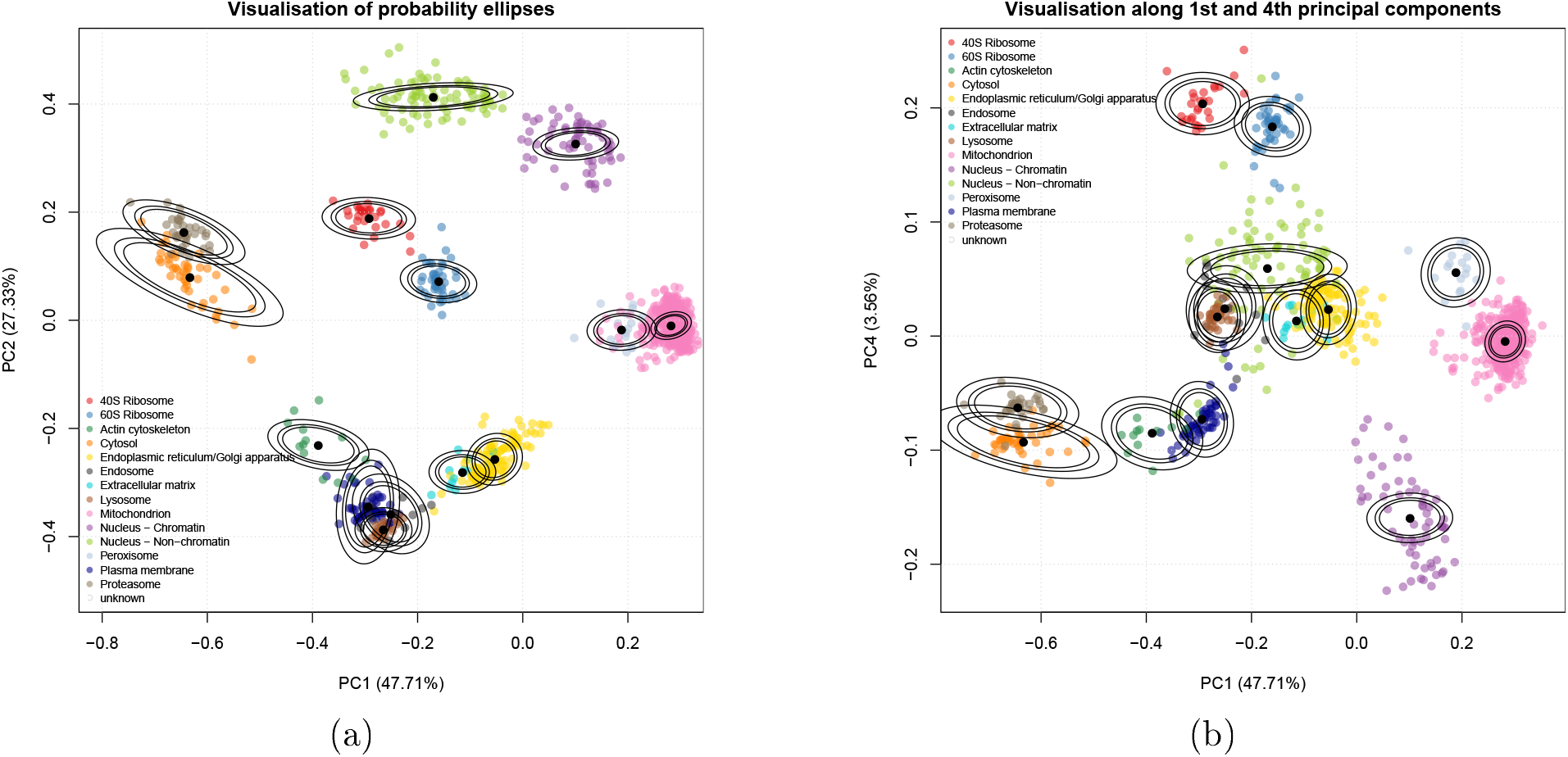
(a) PCA plot of the 1st and 2nd principal components for the curated marker proteins of the mouse stem cell data. The organelles are, in general, well separated. Though some organelles overlap, they are separated along different principal components. The densities used to produce the ellipses are derived from the MAP estimates, (b) Marker resolution along the 1st and 4th principal components show that the mitochondrion and peroxisome markers are well resolved, despite overlapping in the 1st and 2nd component. We also see that the ER/Golgi apparatus markers are better separated from the extracellular matrix markers.

We now apply the statistical methodology described in section 4, to predict the localisation of proteins to organelles and sub-cellular components. In brief, we produce MAP estimates of the parameters by using the expectation-maximisation algorithm, to form the basis of a Bayesian analysis (TAGM-MAP), We run the algorithm for 200 iterations and inspect a plot of the log-posterior to assess convergence of the algorithm (see appendix 5.3), We confirm that the difference of the log posterior between the final two iterations is less than 10^−6^ and we conclude that our algorithm has converged. The results can be seen in figure 2 (left), where the posterior localisation probability is visualised by scaling the pointer for each protein.

Figure 2 (right) demonstrates a range of probabilistic assignments of proteins to organelles and sub-cellular niches. We additionally consider a full, sampling-based Bayesian analysis using Markov-chain Monte Carlo (MCMC) to characterise the uncertainty in the localisation probabilities. In our case a collapsed Gibbs sampler is used to sample from the posterior of localisation probabilities. The remainder of this article focus on analysis of spatial proteomics in this fully Bayesian framework.

**Figure 2:**
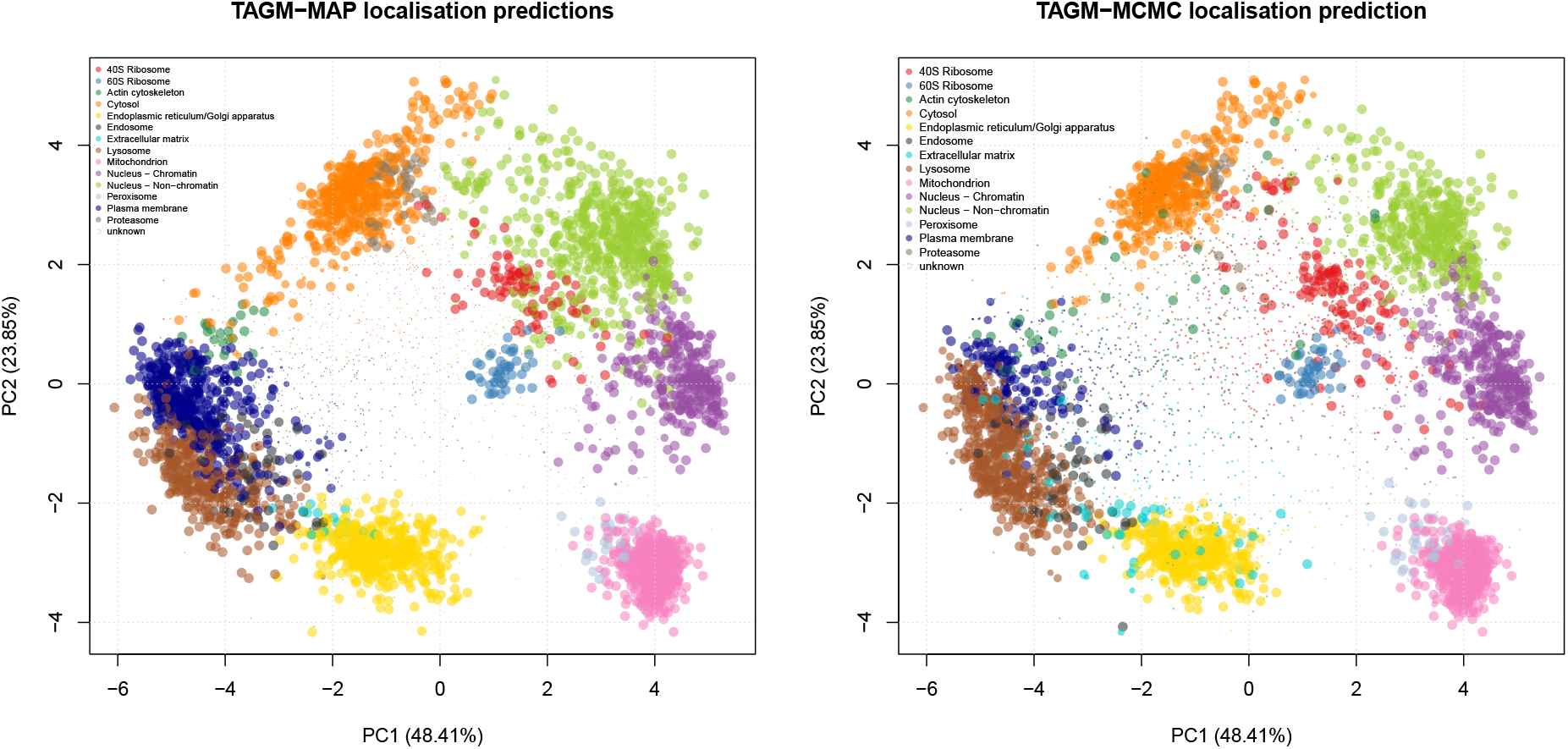
PCA plot of the protein quantitation data with colours representing the predicted class (5032 proteins) illustrating protein localisation preductions using TAGM-MAP (left) and TAGM-MCMC (right) respectively. The pointer size of a protein is scaled to the probability that particular protein was assigned to that organelle. Markers, proteins whose localisations are already known, are automatically assigned a probability of 1 and the size of the pointer reflects this.

#### 2.1.2 Uncertainty in the posterior localisation probabilities

This section applies the TAGM model to the mouse pluripotent embryonic stem cell data, by considering the uncertainty in the parameters and exploring how this uncertainty propogates to the uncertainty in protein localisation prediction. In figure 3 we visualise the model as before using the first two principal components along with the first and fourth principal component as a representive example. For the TAGM model, we derive probability ellipses from the expected value of the posterior normal-inverse-Wishart (NIW) distribution.

We apply the statistical methodology detailed in section 4, We perform posterior computation in the Bayesian setting using standard MCMC methods (TAGM-MCMC), We run 6 chains of our Gibbs sampler in parallel for 15,000 iterations, throwing away the first 4, 000 iterations for burn-in and retain every 10^*th*^ sample for thinning. Thus 1,100 sample are retained from each chain. We then visualise the trace plots of our chains; in particular, we monitor the number of proteins allocated to the known components (see appendix 5.4). We discard 1 chain because we do not consider it to have converged. For the remaining 5 chains 184 we further discard the first 500 samples by visual inspection. We then have 600 retained samples from 5 separate chains. For further analysis, we compute the Gelman-Rubin convergence diagnostic (Gelman and Rubin, 1992; Brooks and Golman, 1998), which is computed as 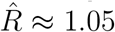. Values of 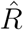 far from 1 indicate non-convergence and since our statistic is less than 1.1, we conclude our chains have converged. The remaining samples are then pooled to produce a single chain containing 3000 samples.

We produce point estimates of the posterior localisation probabilities by summarising samples by their Monte-Carlo average. These summmaries are then visualised in figure 2 (right panel), where the pointer is scaled according to the localisation probabilities of the sub-cellular niche with the largest posterior probability, Monte-Carlo based inference also provides us with additional information; in particular, we can interrogate individual proteins and their posterior probability distribution over sub-cellular locations.

Figure 4 illustrates one example of the importance of capturing uncertainty. The E3 ubiquitin-protein ligase TRIP12 (G5E870) is an integral part of ubiquitin fusion degradation pathway and is a protein of great interest in cancer because it regulates DNA repair pathways. The SVM failed to assign this protein to any location, with assigment to the 60S Ribosome falling below a 5% FDR and the MAP estimate assigned the protein to the nucleus non-chromatin with posterior probability < 0.95. The posterior distribution of localisation probabilities inferred from the TAGM-MCMC model, shown in figure 4, demonstrates that this protein is most probably localised to the nucleus non-chromatin. However, there is some uncertainty about whether it localises to the 40S ribosome. This could suggest a dynamic role for this protein, which could be further explored with a more targeted experiment.

**Figure 3:**
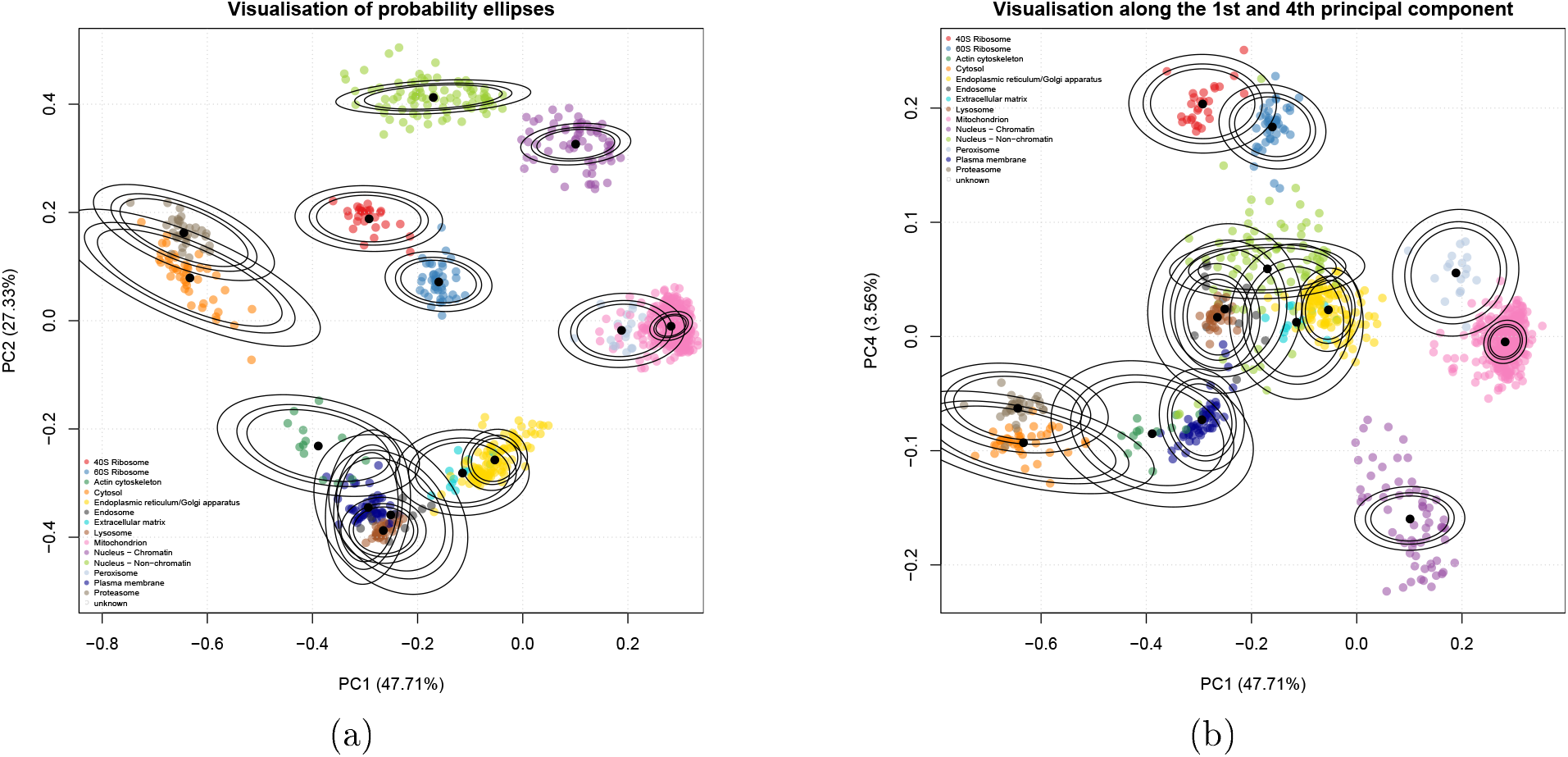
(a) Probability ellipses produced from using the MCMC method. The density is the expected value from the XIW distribution, (b) Probability ellipses visualised along the 1st and 4th principal component also from the MCMC method.

**Figure 4:**
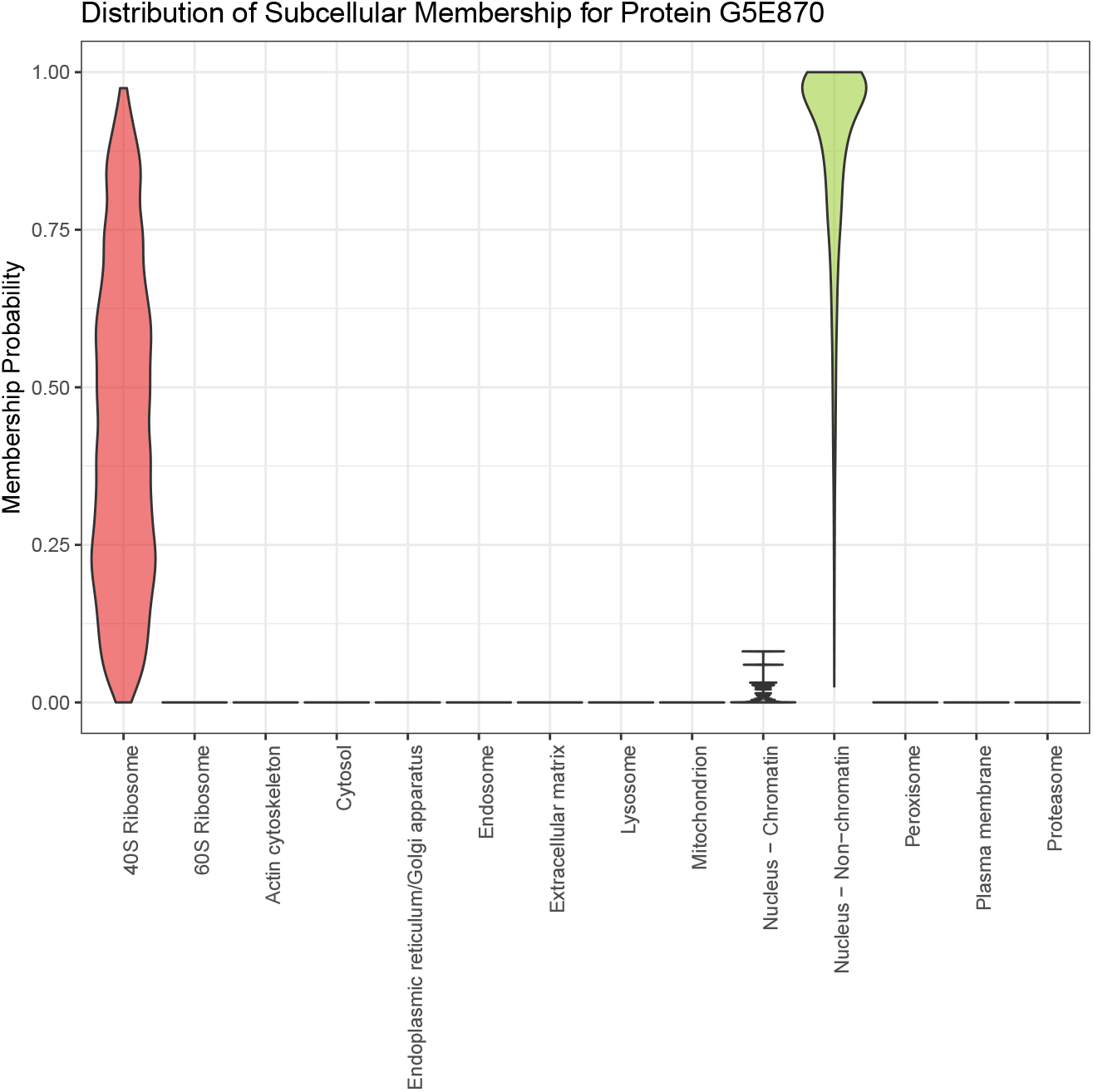
Violin plot revealing the posterior distribution of localisation probabilities of protein E3 ubiquitin-protein ligase (G5E870) to organelles and sub-cellular niches. The most probable localisation is nucleus non-chromatin. however there is uncertainty associated with this assignment.

#### 2.1.3 Enrichment analysis of outlier proteins

In previous sections, we demonstrated that we can assign proteins probabilitically to sub-cellular compartment and quantify the uncertainty in these assignments. Some proteins cannot be well described as belonging to any known component and we model this using an additional T-distribution outlier component (see Section 4).

It is biologically interesting to decipher what functional role proteins that are far away from known components play. We perform an over-representation analysis of gene ontology (GO) terms to asses the biological relevance of the outlier component (Boyle *et al.*, 2004; Yu *et al.*, 2012), We take 1111 proteins that were allocated to known components with probability less than 0.95. Note that these 1111 proteins exclude proteins that are likely to belong to a known location, but we are uncertain about which localisation. We then perform enrichment analysis against the set of all proteins quantified in the *hyshows this outlier componentper*LOPIT experiment. We search against the cellular compartment, biological process and molecular function ontologies.

Supplementary figure 16 shows this outlier component is enriched for cytoskeletal part (*p* < 10^−7^) and microtuble cvtoskeleton (*p* < 10^−7^), Cytoskeleton proteins are found throughout the cell and therefore we would expect them to be found in every fraction along the density gradient. We also observe enrichment for highly dynamic sub-cellular processes such as cell division (*p* < 10^−6^) and cell cycle processes (*p* < 10^−6^), again these proteins are unlikely to have steady-state locations within a single component. We also see enrichment for molecular functions such as tranferase activity (*p* < 0.005), another highly dynamic process. These observations justify including an additional outlier component in our mixture model.

### 2.2 Comparison with other classifiers

In this section, we assess the generalisation performance of our methods on several datasets, by computing performance metrics associated with each classifier as detailed in section 4.4, We compare the SVM and KNN classifiers alongside the MAP and MCMC approaches detailed in the methods section. We compute the FI score and quadratic loss over 100 rounds of stratified 5-fold cross-validation. The hyperparameter for the KNN algorithm, the number of nearest neighbours, is optimised via an additional internal 5-fold cross-validation and the hyperparameters for the SVM, sigma and cost, are also optimised via internal 5-fold cross validation (Hsu *et al.*, 2010).

We test our methods on the following datasets *Drosophila* (Tan *et al.*, 2009), chicken (Hall *et al.*, 2009), mouse pluripotent embryonic stem cells from Christoforou *et al.* (2016) and Breckels *et al.* (2016b), the human bone osteosarcoma epithelial (U2-OS) cell line (Thul *et al.*, 2017), the HeLa cell line of Itzhak et al. (2016), the 3 HeLa cell lines from Hirst *et al.* (2018) and 10 primary fibroblast datasets from Beltran *et al.* (2016). These datasets represent a great variety of spatial proteomics experiments across many different workflows. The two *hyper*LOPIT datasets on mouse pluripotent embryonic stem cells and the U2-OS cell line use TMT 10-plex labelling and contain the greatest number of proteins. Earlier LOPIT experiments on the *Drosophila* and chicken use iTRAQ 4-plex labelling, whilst another LOPIT mouse pluripotent embryonic stem cell dataset uses iTRAQ 8-plex, The datasets of Itzhak *et al.* (2016) and Hirst *et al.* (2018) employ a different methodology completely - seperating cellular content using differential centrifugation (as opposed to along a density-gradient). Furthermore, the methods use SILAC rather than iTRAQ or TMT for labelling. The experiments of Hirst *et al.* (2018) were designed to explore the functional role of AP-5 by coupling CRISPR-CAS9 knockouts with spatial proteomics methods. We analysed all three datasets from Hirst *et al.* (2018), which includes a wild type HeLa cell line as a control, as well as two CRISPR-CAS9 knockouts: AP5Z1-K01 and AP5Z1-K02 respectively.

In addition, we analyse the spatio-temporal proteomics experiments of Beltran *et al.* (2016), which uses TMT-based MS quantification. This experiment explored infecting primary fibroblasts with Human cytomegalovirus (HMCV) and the goal of these experiments was to explore the dynamic perturbation of host proteins during infection, as well as the sub-cellular localisation of viral proteins throught the HCMV life-cycle. They produced spatial maps at different time points: 24, 48, 72, 96, 120 hours post infection (hpi), as well as mock maps at these same time points to serve as a control - this results in 10 different spatial proteomics maps.

In each case, a dataset specific marker list was used, which is curated specifically for the each cell line. We removed “high-curvature ER” annotations from the HeLa dataset (Itzhak *et al.*, 2016), as well as the “ER Tubular”, “Nuclear pore complex” and “Peroxisome” annotations from the HeLa CRISPR-CAS9 knockout experiments (Hirst *et al.*, 2018) as there are too few proteins to correctly perform cross-validation. Table 1 summarises these datasets, including information about number of quantified proteins, the workflow used and the number of fractions.

**Table 1:** Summary of spatial proteomics datasets used for comparisons

| MS-based Spatial Proteomics datasets |  |  |  |  |
| --- | --- | --- | --- | --- |
| Cell line or organism | Workflow | Labelling | Fractions (including combined replicates) | Proteins |
| *Drosophila* | LOPIT | iTRAQ | 4 | 888 |
| Chicken DT40 | LOPIT | iTRAQ | 16 | 1090 |
| Mouse pluripotent E14TG2a stem cell | HyperLOPIT | TMT | 20 | 5032 |
| HeLa (Itzhak et al.) | Organeller Maps | SILAC | 30 | 3766 |
| HeLa (Hirst et al.) | Organeller Maps | SILAC | 15 | 2046 |
| U2-OS cell line | HyperLOPIT | TMT | 37 | 5020 |
| Primary Fibroblast | Spatio-Temporal Methods | TMT | 6 | 2196 |
| E14TG2a (Breckels et al.) | LOPIT | iTRAQ | 8 | 2031 |

Figure 5 compares the Macro-F1 scores across the datasets for all classifiers and demonstrates that no single classifier consistently outperforms any other across all datasets, with results being highly consistent across all methods, as well as across datasets. We perform a pairwise unpaired t-test with multiple testing correction applied using the Benjamini-Höchberg procedure (Benjamini and Hochberg, 1995) to detect differences between classifier performance.

In the *Drosophila* dataset only the KNN algorithm outpeforms the SVM at significance level of 0.01, whilst no other significant differences exist between the classifiers. In the chicken DT40 dataset only the MCMC method outperforms the KNN classifier at significance level of 0.01, no other significant conclusion can be drawn. In the mouse dataset the MAP based method outperforms the MCMC method at significance level of 0.01, no other significant conclusions can be drawn. In the HeLa dataset all classifiers are significantly different at a 0.01 level. These differences may exist because the dataset does not fit well with our modelling assumptions; in particular, this dataset set has been curated to have a class called “Large Protein Complex”, which likely describes several sub-cellular structures. These might include nuclear compartments and ribosomes, as well as any cytosolic complex and large protein complex which pellets during the centrifugation conditions used to capture this mixed sub-cellular fraction. Moreover, the cytosolic and nuclear fraction were processed separately leading to possible imbalance with comparisons with other datasets. Thus, the large protein complex component might be better described as itself a mixture model or more detailed curation of these data may be required. We do not consider further modelling of this dataset in this manuscript. For the U2-OS all classifiers are significantly different at a significance level of 0.01 except for the SVM classifier and the MCMC method, with the MAP method performing the best. Figure 5 shows that for this dataset all classifiers are performing extremely well. In the three Hirst datasets the MAP method significantly outperforms all other methods (*p* < 0.01), whilst in the wild type HeLa and in the CRISPR-CAS9 KO1 there is no significant difference between the KNN and MCMC method. In the CRISPR-CAS9 KO2 the MCMC method outperforms the SVM and KNN methods (*p* < 0.01), In the interest of brevity, the remaining results for the t-tests can be found in tables in appendix 5.5.

**Figure 5:**
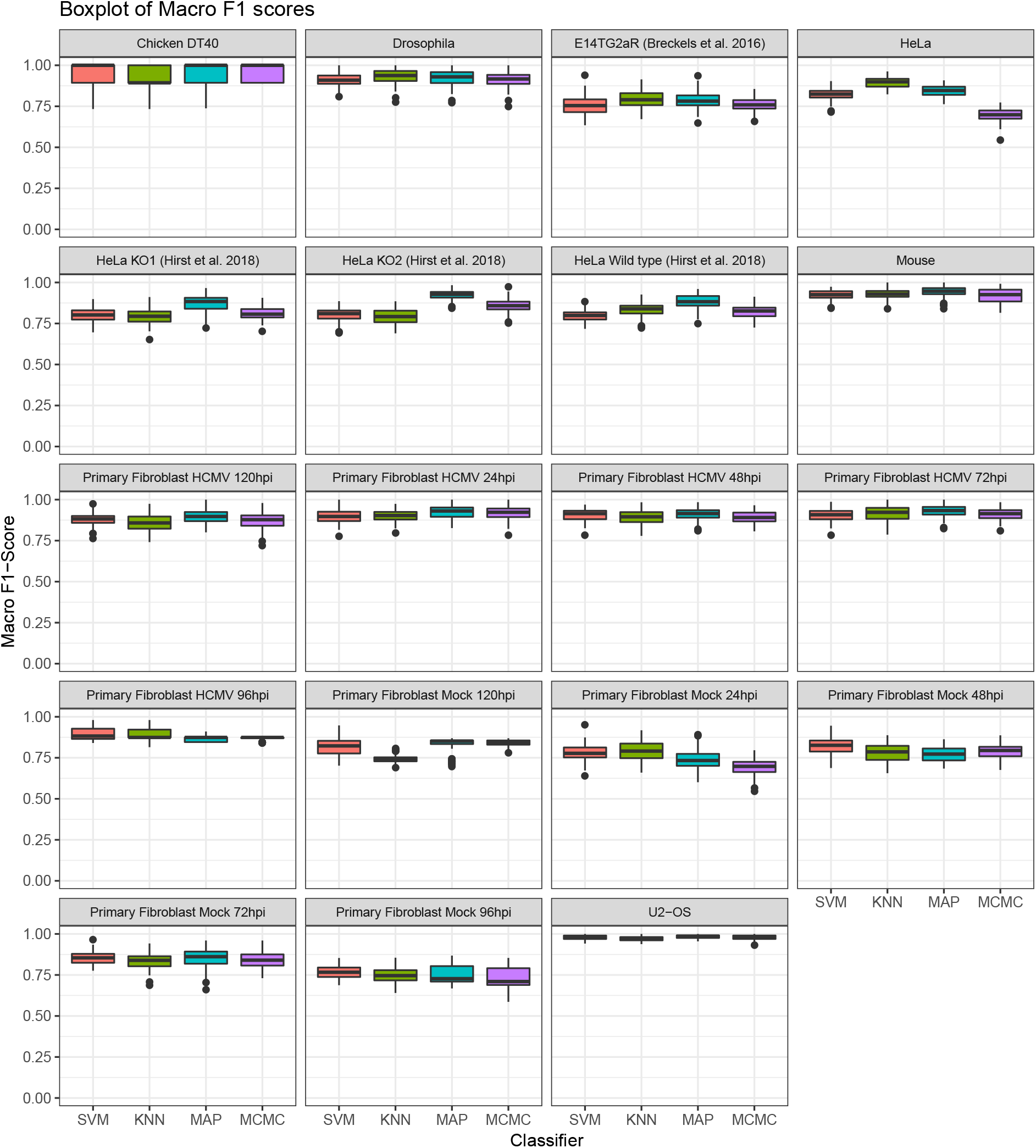
Boxplots of the distributions of Macro F1 scores for all spatial proteomics datasets.

The Macro-F1 scores do not take into account that whilst the TAGM model may mis-classify, it may do so with low confidence. We therefore additionally compute the quadratic loss, which allows us to make use of the probabilitic information provided by the classifiers. The lower the quadratic loss the closer the probabilitic predicition is to the true value. We plot the distributions of quadratic losses for each classifier in figure 6, We observe highly consitent performance across all classifiers across all datasets. Again, we perform a pairwise unpaired t-test with multiple testing correction.

We find that in 16 out of 19 datasets (all of those except HeLa Wild type, HeLa KOI and HeLa KO2) the MCMC methods achieves the lowest quadratic loss at a signifiance level < 0.0001 over the SVM and KNN classifiers. In 6 out of these 16 datasets there is no significant difference between the MCMC and the MAP methods. In the three Hirst datasets in which the MCMC did not acheive the lowest quadratic loss, the SVM outperformed. However, in two of these datasets (HeLa Wild type and KOI) the MAP method and SVM classifier were not significantly different. In the Hirst KO2 dataset there were no signicant differences between the MAP and MCMC methods.

In the vast majority of cases, we observe that if the TAGM model, using the MCMC methodology, makes an incorrect classification it does so with lower confidence than the SVM classifier, the KNN classifier and the MAP based classifier, whilst if it is correct in its assertion it does so with greater confidence. Additionally, a fully Bayesian methodology provides us with not only point estimates of classification probabilities but uncertainty quantification in these allocations, and we show in the following section that this provides deeper insights into protein localisation.

**Figure 6:**
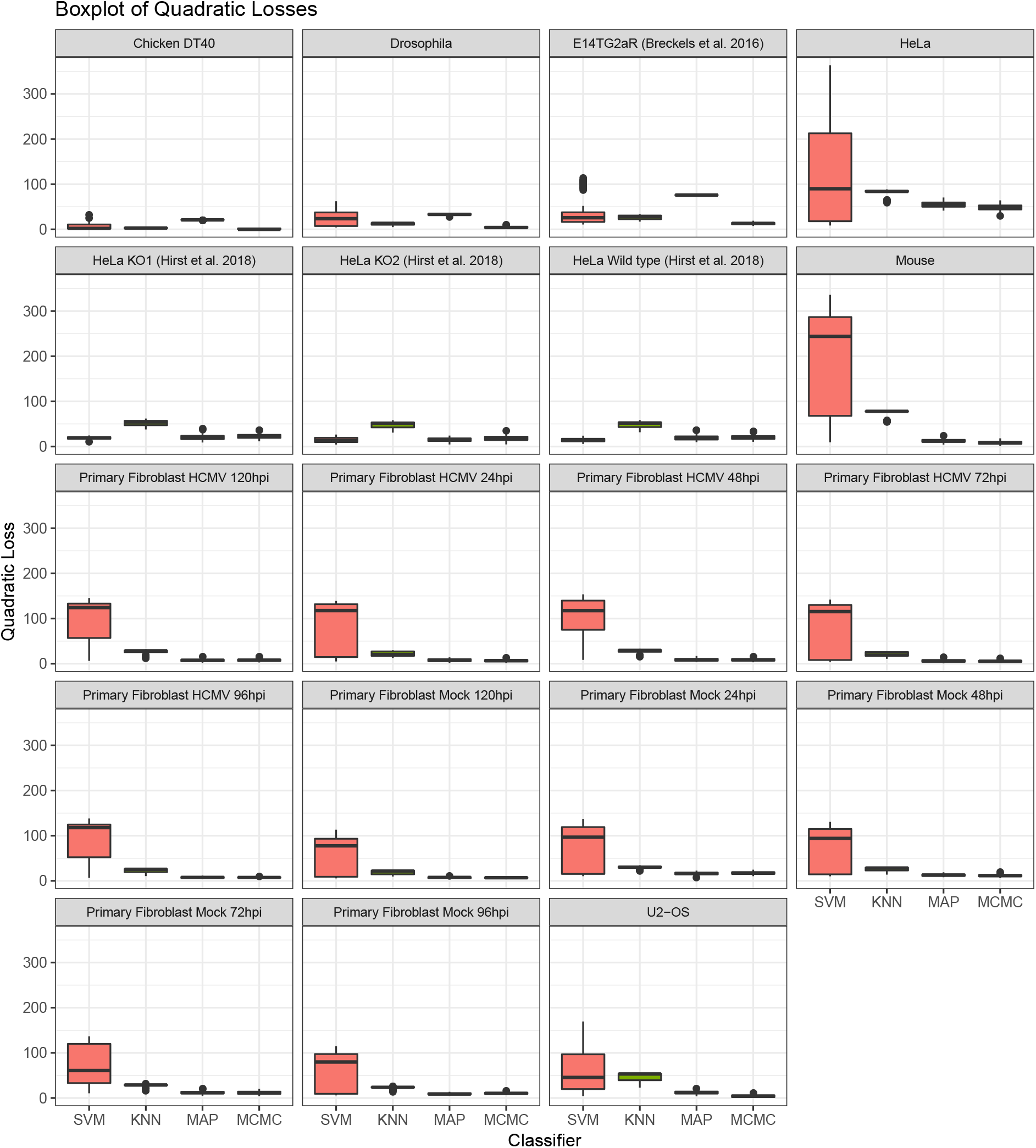
Boxplots of the distributions of Quadratic losses for all spatial proteomics datasets.

Computing distributions of FI scores and quadratic losses, which can only be done on the marker proteins, can help us understand whether a classifier might have greater generalised performance accuracy. However, we are interested in whether there is a large disagreement between classifiers when prediction is performed on proteins for which we have no withheld localisation information. This informs us about a systematic bias for a particular classifier or whether a classifier ensemble could increase performance. To maintain a common set of proteins we set thresholds for each classifier in turn and compare to the other classifier without thresholding. Firstly, we set a global threshold of 0,95 for the TAGM-MCMC and then for these proteins plot a contingency table against the classification results from the SVM. Secondly, we set a *5%* FDR for the SVM and then for these proteins plot a contingency table against the classification results from the TAGM-MCMC, We visualise the contingency tables as heat plots in figure 7.

**Figure 7:**
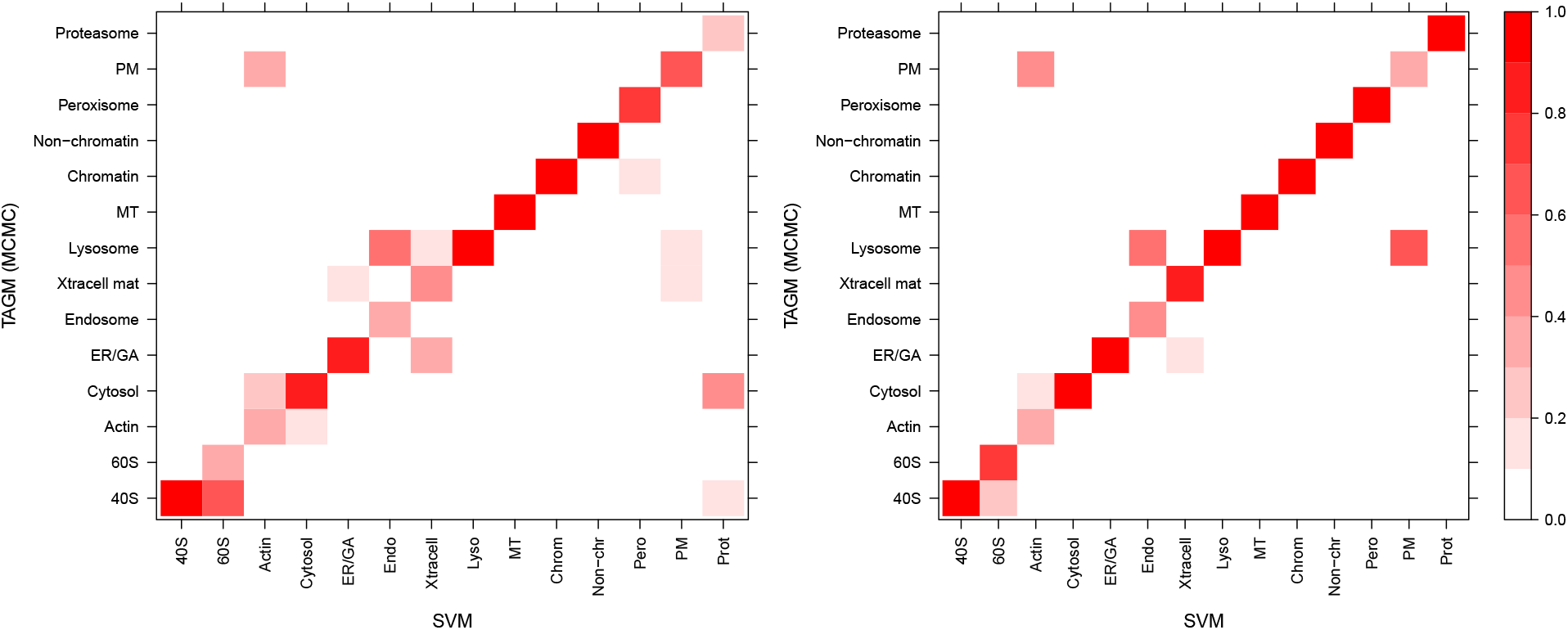
A heatmap representation of a contingency table, where we compare assignment results for proteins with unknown protein localisation using the TAGM-MCMC and SVM, The scale ranges from 0 to 1 with values indicating the proportion of assigned proteins to that sub-cellular location. Values along the diagonal represent agreement between classifiers whilst other values represent disagreement. The coherence between the classifers is very high, (a) In this case we set a probability threshold of 0,95 for the TAGM assignments with no threshold for the SVM, (b) In this case we set a *5%* FDR threshold for the SVM and no threshold for the TAGM-MCMC.

In general, we see an extremely high level of coherence between the TAGM and the SVM, with almost all proteins predicted to concordant sub-cellular compartments. Figure 7 shows there is some disagreement between assigning proteins to the lysosome and plasma membrane, to the cytosol and proteasome, and between the large and small ribosomal subunits. However, we have not used the uncertainty in the probabilitic assignments to produce the contingency tables above. In the next sections, we explore examples of proteins with uncertainty in their posterior localisation probabilities. Selecting biologically relevant thresholds is important for any classifier and exploring uncertainty is of vital importance when drawing biological conclusions.

### 2.3 Interpreting and exploring uncertainty

Protein sub-cellular localisation can be uncertain for a number of reasons. Technical variations and unknown biological novelty, such as yet uncharacterised functional compartments, can be some of the reasons why a protein might have an unknown or uncertain localisation. Furthermore many proteins are known to reside in multiple locations with possibly different functional duties in each location (Jeffery, 2009), With these considerations in mind, it is pertinent to quantify the uncertainty in our allocation of proteins to organelles. This section explores several situations where proteins display uncertain localisation and considers the biological factors that influence uncertainty. We later explore and visualise whole proteome uncertainty quantification.

Exportin 5 (Q924C1) forms part of the micro-RNA export machinery of the nucleus, transporting miRNA from the nucleus to the cytoplasm for further processing. It then translocates back through the nuclear pore complex to return to the nucleus, Exportin 5 can then continue to mediate further transport between nucleus and cytoplasm. The SVM was unable to assign a localisation of Exportin 5, with its assignment falling below a 5% FDR to wrongly assign this protein to the proteasome. This incorrect assertion by the SVM was confounded by the similarity between the cytosol and proteasome profiles. Figure 8 demonstrates, according to the TAGM-MCMC model, that Exportin 5 most likely localises to the cytosol but there is some uncertainty with this assignment. This uncertainty is reflected in possible assignment of Exportin 5 to the nucleus non-chromatin and this uncertainty is a manifestation of the the fact that the function of this protein is to shuttle between the cytosol and nucleus.

The Phenylalanine-tRNA ligase beta subunit protein (Q9WUA2) has an uncertain localisation between the 40S ribosome and the nucleus non-chromatin demonstrated in figure 9. This protein was left unclassified by the SVM because its score fell below a threshold to assign it to the 40S ribosome. Considering that this protein is involved in the acylation of transfer RNA (tRNA) with the amino acid phenylalanine to form tRNA–Phe to be used in translation of proteins, it is therefore unsurprising that this protein’ s steady state location is ribosomal. Whilst the SVM is unable to make an assignment, TAGM-MCMC is able to suggest an assignment and quantify our uncertainty.

Relatively little is known about the Dedicator of cytokinesis (DOCK) protein 6 (Q8VDR9), a guanine nucleotide exchange factor for CDC42 and RAC1 small GTPases, The SVM could not assign localisation to the ER/Golgi, since its score fell below a 5% FDR. Furthermore, the TAGM-MCMC model assigned this DOCK 6 to the outlier component with posterior probability > 0.95. Figure shows possible localisation to several components along the secretory pathway. As an activator for CDC42 and RAC1 we may expect to see them with similar localisation, CDC42, a plasma membrane associated protein, regulates cell cycle and division and is found with many localisations. Furthermore RAC1, a small GTPase, also regulates many cellular processes and is found in many locations. Thus the steady-state distribution of DOCK6 is unlikely to be in a single location, since its interaction partners are found in many locations. This justifies including an outlier component in our model, else we may erroneously assign such proteins to a single location.

**Figure 8:**
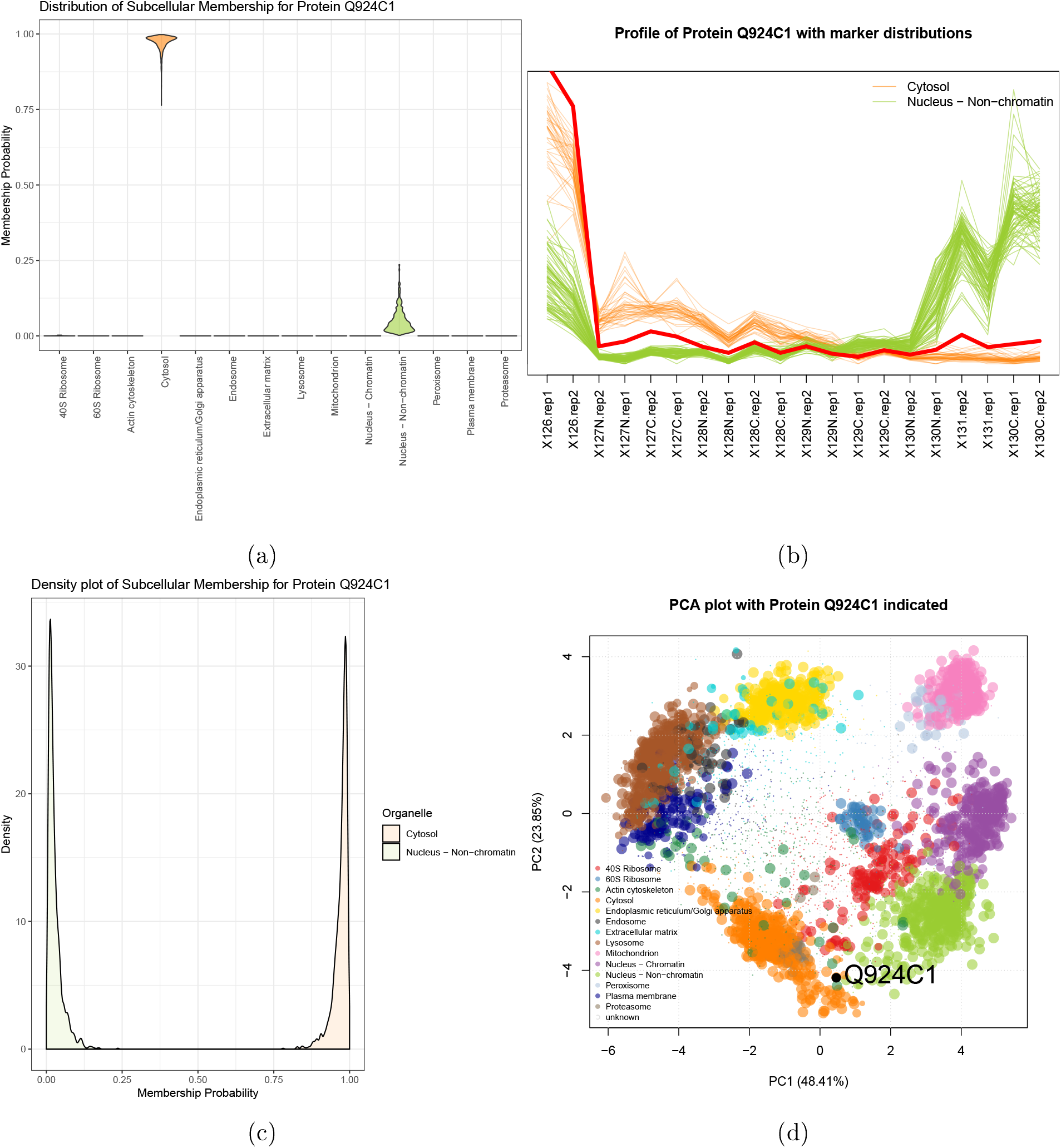
Export in 5 (Q924C1) showing localisation to the cytosol with some uncertainty about association to the nucleus non-chromatin. (a) The violin plot shows uncertain localisation between these two sub-cellular localisations. (b) The quantitative profile of this protein shows mixed profile between the profiles of the organelle markers. (c) The density plot shows a complex distribution over localisations for this protein. (d) The protein Q924C1 has steady state distribution between the cytosol and nucleus non-chromatin.

**Figure 9:**
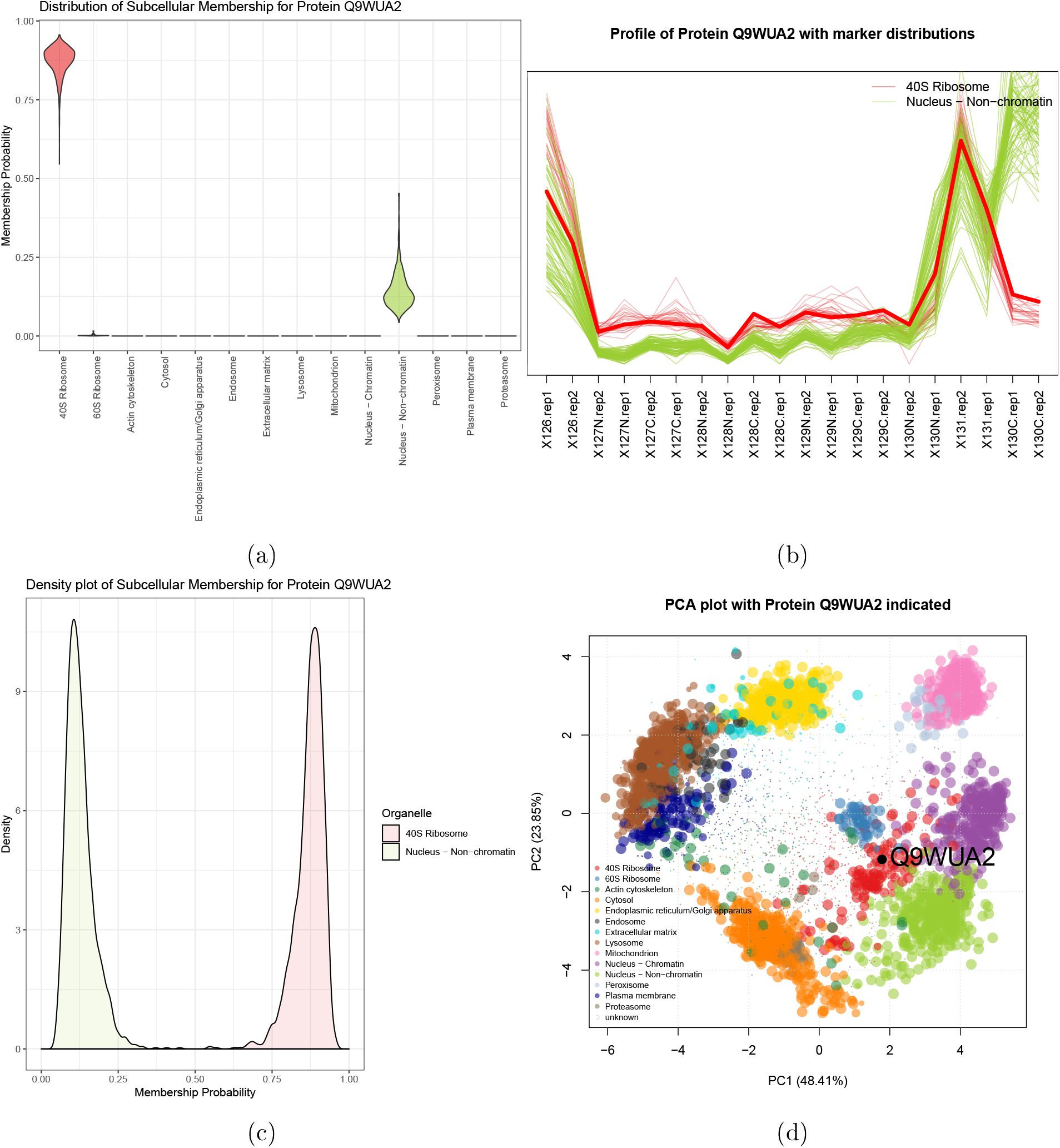
Phenylalanine-tRNA ligase beta subunit protein TRIP12 (Q9WUA2) showing localisation to the 40S Ribosome with some uncertainty about association to the nucleus non-chromatin, (a) The violin plot shows uncertain localisation between these two sub-cellular localisations, (b) The quantitative profile of this protein shows mixed profile between the profiles of the organelle markers. (c) The density plot shows a complex distribution over localisations for this protein. (d) The protein Q9WUA2 has steady state distribution skewed towards the 40S Ribosome and close to the nucleus non-chromatin.

**Figure 10:**
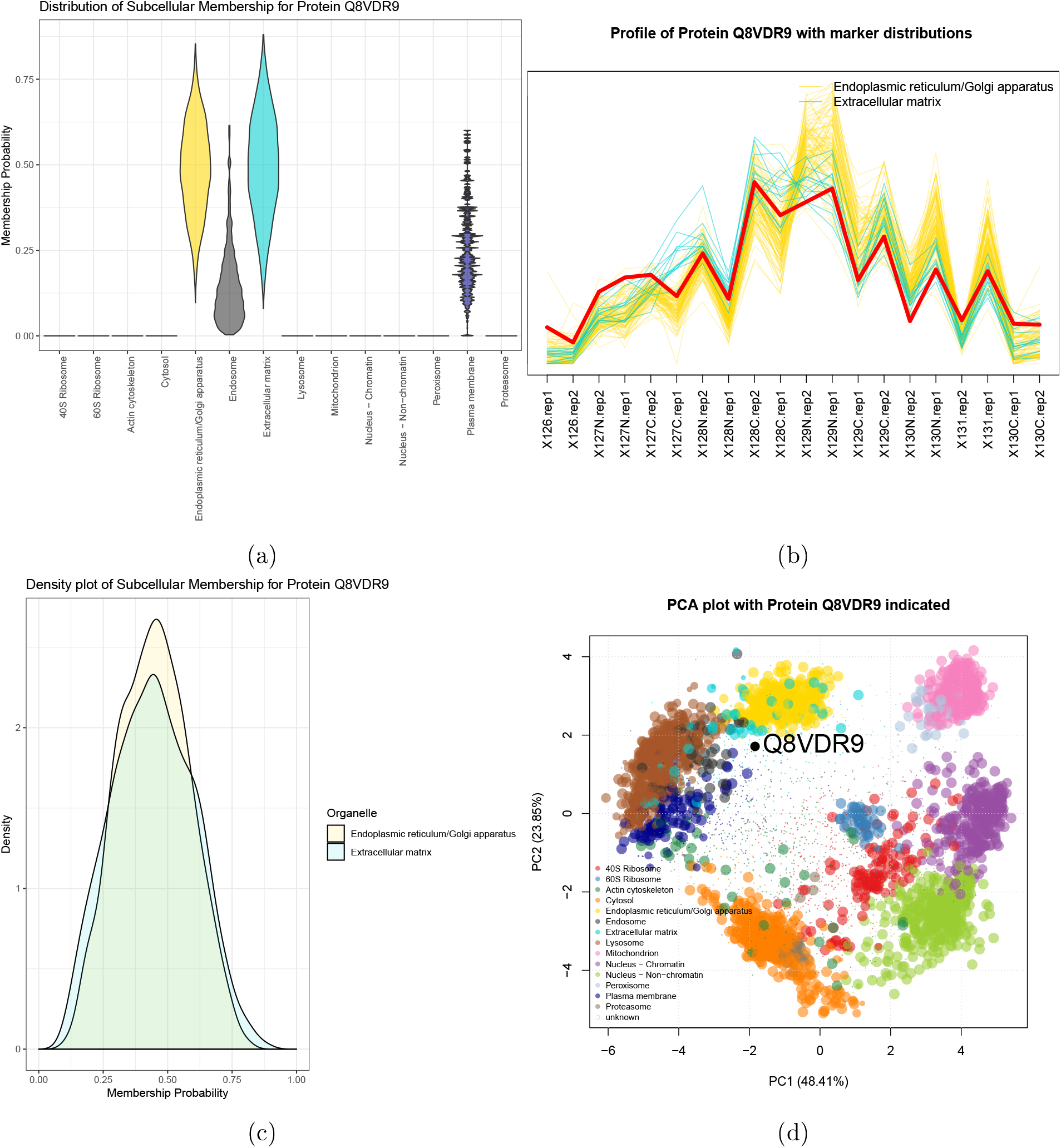
Q8VDR9 showing localisation to the outlier component. (a) The violin plot shows uncertain localisation between several sub-cellular niches. (b) The quantitative profile of this protein shows mixed profile between the profiles of the organelle markers. (c) The density plot shows a similar localisation probabilities for both the ER/Golgi and Extracellular matrix. (d) The protein Q8VDR9 has steady state distribution in the centre of the plot skewed toward the secretory pathway; in particular, the ER/Golgi and Extracellular matrix components.

### 2.4 Visualising whole sub-cellular proteome uncertainty

The advantage of the TAGM-MCMC model is its ability to provide proteome wide uncertainy quantification. Regions where organelle assignments overlap are areas were uncertainty is expected to be the greatest, as well as areas with no dominant component. We take an information theoretic approach to summarising uncertainty in protein localisation by computing the Shannon entropy (Shannon, 1948) for each Monte-Carlo sample *t* = 1, …, *T* of the posterior localisation probabilities of each protein

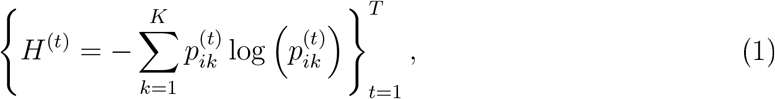

where 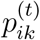 denotes the posterior localisation probabiltv of protein *i* to component *k* at iteration *t*. We then summarise this as a Monte-Carlo averaged Shannon entropy. The greater the Shannon entropy the more uncertainty associated with the assignment of this protein. The lower the Shannon entropy the lower the uncertainty associated with the assignment of this protein. In figure 11 panel (a), we visualise the Shannon entropy of each protein in a PCA plot, by scaling the pointers in accordance to this metric. We also note that while localisation probability (of a protein to its most probable location) and the Shannon entropy are correlated, figure 11 panel (c), it is not perfect. Thus it is important to use both the localisation probabilities and the uncertainty in these assignments to make conclusions.

Figure 11 demonstrates that the regions of highest uncertainty are those in regions where organelles assignments overlap. The conclusions from this plot are manifold. Firstly, many proteins are assigned unambiguously to sub-cellular localisations; that is, not only are some proteins assigned to organelles with high probability but also with low uncertainty. Secondly, there are well defined regions with high uncertainty, for example proteins in the secretory pathway or proteins on the boundary between cytosol and proteasome. Finally, some organelles, such as the mitochondria, are extremely well resolved. This observed uncertainty in the secretory pathway and cytosol could be attributed to the dynamic nature of these parts of the cell with numerous examples of proteins that traffic in and out of these sub-cellular compartments as part of their biological role. Moreover, the organelles of the secretory pathway share similar and overlapping physical properties making their separation from one another using biochemical fractionation more challenging. Furthermore, there is a region located in the centre of the plot where proteins simultaneously have low probability of belonging to any organelle and high uncertainty in their localisation probability. This suggests that these proteins are poorly described by any single location. These proteins could belong to multiple locations or belong to undescribed sub-cellular compartments. The information displayed in these plots and the conclusion therein would be extremely challenging to obtain without the use of Bayesian methodology.

**Figure 11:**
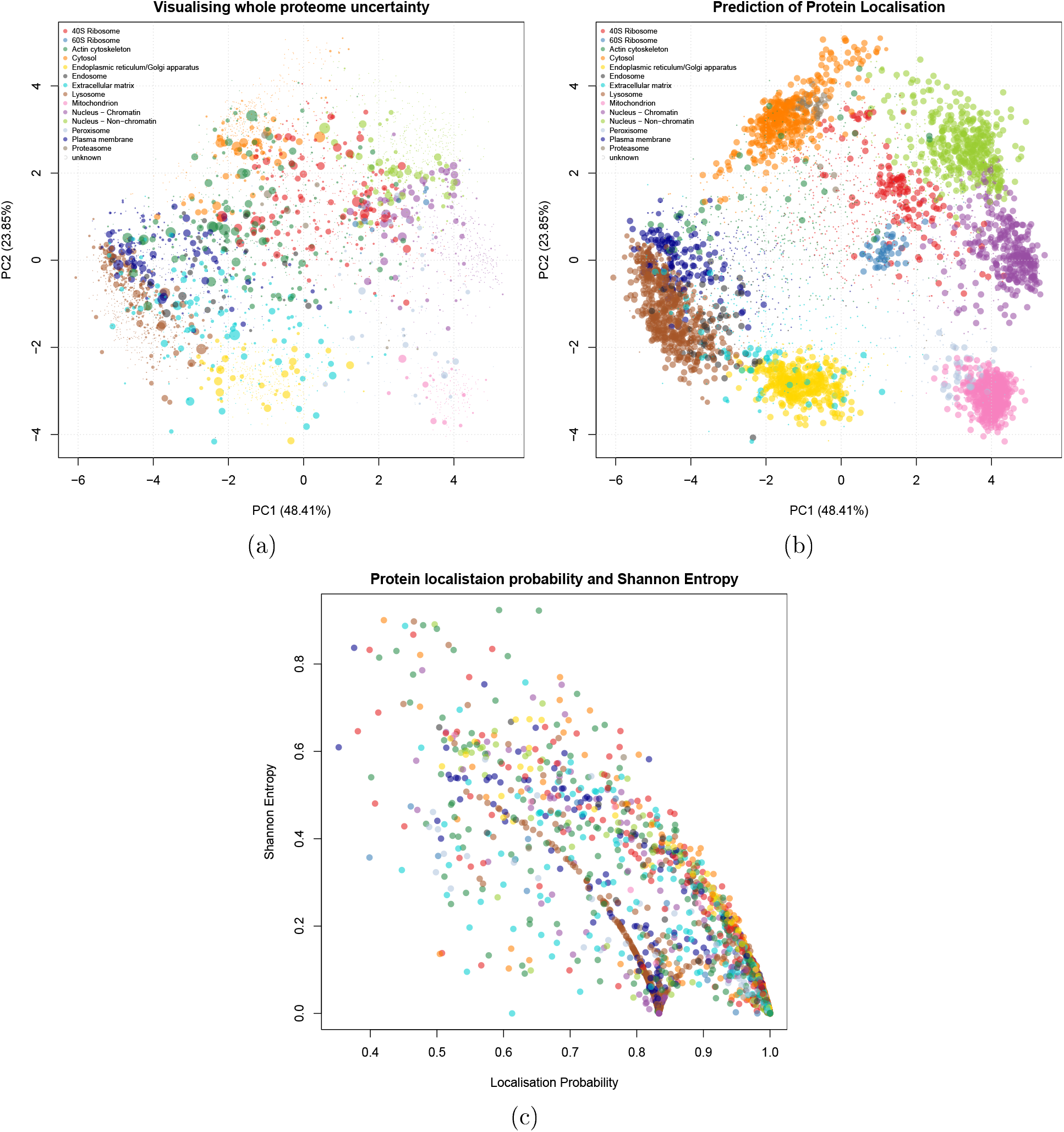
PC A plots of the mouse pluripotent embryonic stem cell data, where each point represents a protein and is coloured to its (probabilistically-)assigned organelle. (a) In this plot, the pointer is scaled to the Shannon entropy of this protein, with larger pointers indicating greater uncertainty. (b) In this plot, the pointer is scaled to the probability of that protein belonging to its assigned organelle. (c) We plot the localisation probabilities against the Shannon entropy with each protein.

## 3 Discussion

We have demonstrated that a Bayesian framework, based on Gaussian mixture models, for spatial proteomics can provide whole sub-cellular proteome uncertainty quantification on the assignment of proteins to organelles and such information is invaluable. Performing MAP inference using our generative model provides fast and straightforward approach, which is vital for quality control and early data exploration. Full posterior inference using MCMC provides not only point estimates of the posterior probability that a protein belongs to a particular sub-cellular niche, but uncertainty in this assignment. Then, this uncertainty can be summarised in several ways, including, but not limited to, equi-tailed credible intervals of the Monte-Carlo samples of posterior localisation probabilities. Posterior distributions for indivdual proteins can then be rigorously interrogated to shed light on their biological mechanisms; such as, transport, signalling and interactions.

As well as the local uncertainty seen by exploring individual proteins, we further explored using a Monte-Carlo averaged Shannon entropy to visualise global uncertainty. Regions of high uncertainty, as measured using this Shannon entropy, reflect highly dynamics regions of the sub-cellular environment. Hence, biologists can now explore uncertainty at different levels and then are able to make quantifiable conclusions and insights about their data. Furthermore, our Bayesian model is interpretable and our inferences are fully conditional on our data, allowing them to be easily modified with changing experimental design.

In addition, we produced competitive classifier performance to the state-of-the-art classifiers, We considered two traditional machine-learning methods: the SVM and KNN classifiers; as well as two classifiers based on our model: a MAP classifier and classification based on MCMC. We compared all methods on 19 different spatial proteomics datasets, across four different organisms. When considering the macro-FI score as a performance metric, no single classifier outperformed another across all datasets. However, using MCMC based inference our method significantly outperforms the SVM and KNN classifiers with respect to the quadratic loss in 16 our of 19 datasets. This allows us to have greater confidence in our conclusions when they are draw from our Bayesian inferences. Furthermore, using MCMC provides a wealth of additional information, and so becomes the method of choice for analysing spatial proteomics data.

Analysis of a *hyper*LOPIT experiment applied to mouse pluripotent embryonic stem cells demonstrated that the additional layer of information that our model provides is biologically relevant and provides further avenues for additional exploration. Moreover, applying our method to a biologically significant dataset now provides the scientific community with localisation information on up to 4000 proteins for the mouse pluripotent stem cell proteome. Figure 12 demonstrates that from an initial input of roughly 1000 marker proteins with *a priori* known location and 4000 unknown proteins with unknown location, SVM and TAGM-MCMC can provide rigorous localisation information on roughly 2000 proteins. However, our methodology, by also considering uncertainty, allows us to obtain information on another 1000 proteins. Thus, we have augmented this dataset by providing uncertainty quantification on the localisation of proteins to their sub-cellular niches, which had been previously unavailable. We note that our method is general enough to be applied to many MS-based spatial proteomics protocols including: LOPIT, *hyper*LOPIT, protein correlation profiling (PCP) (Foster *et al.*, 2006), differential centrifugation approaches and spatio-temporal proteomics methods. In our flexible software implementation, all hyperparameters for the priors can be changed if users have precise priors they wish to specify.

**Figure 12:**
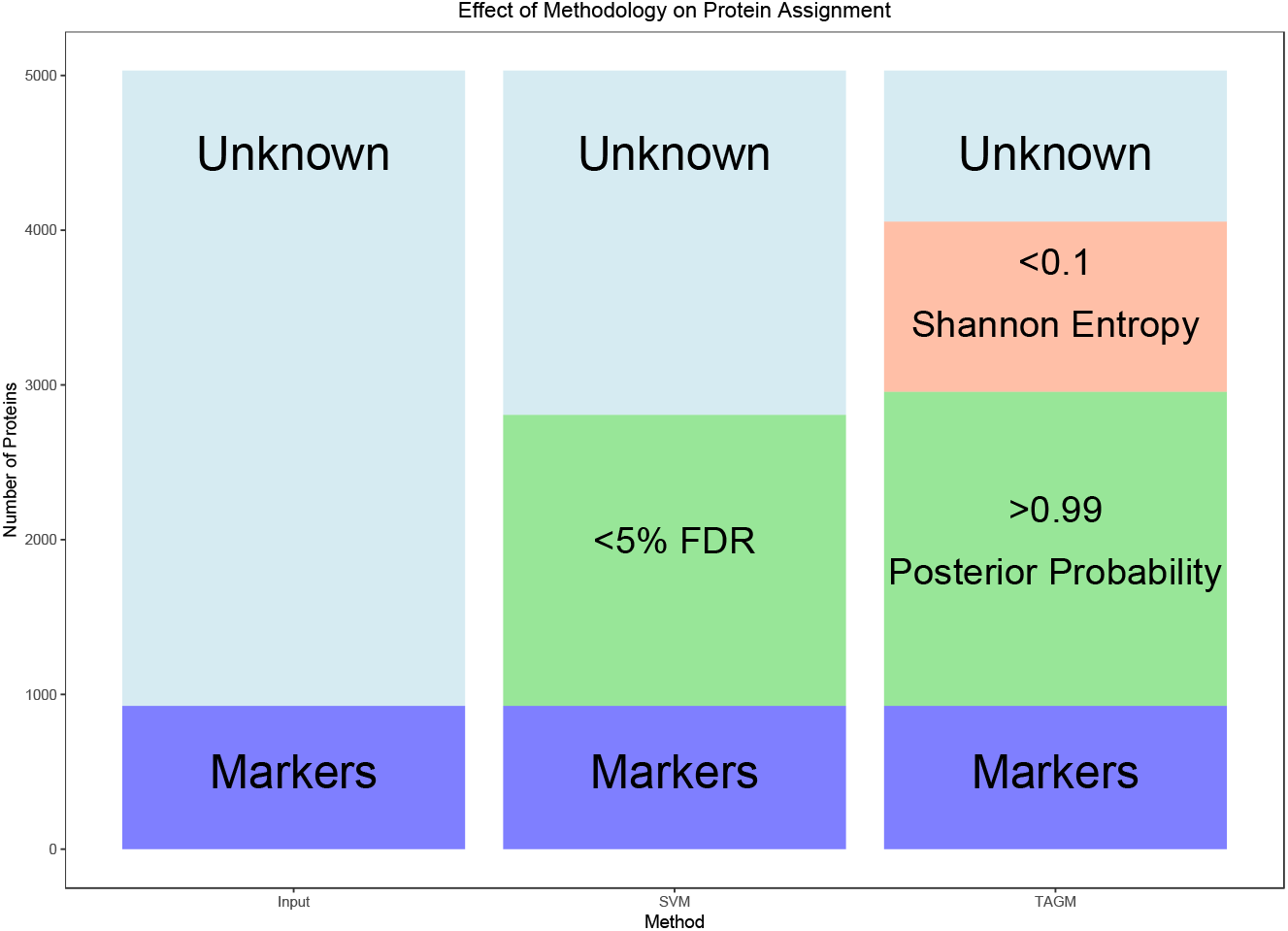
The barplot demonstrates the effect of applying different methodologies on protein assignment when applied the mouse pluripotent embryonic stem cell data. Roughly 2000 proteins are classified using either SVM and TAGM-MCMC; however, TAGM-MCMC can draw additional conclusions about an extra 1000 proteins by quantifying uncertainty.

We have also provided a new set of visualisation methods to accompany our model, which allow us to easily interrogate our data. High quality visualisation tools are essential for rigorous quality control and sound biological conclusions. Our methods have been developed in the R statistical programming language and we continue to contribute to the Bioconductor project (Gentleman *et al.*, 2004; Huber *et al.*, 2015) with inclusion of our methods within the pRoloc package (>= 1.21.1) (Gatto *et al.*, 2014b), The underlying source code used to generate this document is available at https://github.com/lgatto/2018-TAGM-paper.

Currently, our model does not integrate localisation information from different data sources, nor does it explicitly model proteins with multiple localisation. However, one (of many) biological explanations for the uncertainty that we model in the allocation probabilities is provided by multiple localisation. Thus a protein for which it is uncertain to which two sub-cellular niches it is resident within it is perhaps resident of both niches. In further work, we plan to explicitly model such cases to deconvolute different sources of uncertainty. In addition, extensions to semi-supervised non-parametric methods are under consideration to detect novel sub-cellular niches. These are the subjects of further work.

## 4 Model and methods

We describe in this section the probabilistic model that uses the labelled data to associate un-annotated proteins to specific organelles or sub-cellular compartments.

### 4.1 Mixture models for spatial proteomic data

We observe *N* protein profiles each of length *L*, corresponding to the number of quantified fractions along the gradient density, including combining replicates. For *i* = 1, …, *N*, we denote the profile of the i-th protein by x_*i*_ = [*x*_1*i*_, …, *x_L_i*], We suppose that there are *K* known sub-cellular compartments to which each protein could localise (e.g. cytoplasm, endoplasmic reticulum, mitochondria, …), Henceforth, we refer to these K sub-cellular compartments as *components*, and introduce component labels *z_i_*, so that *z_i_* = *k* if the *i*-th protein localises to the *k*-th component. We denote by *X_L_* the set of proteins whose component labels are known, and by *X_U_* the set of unlabelled proteins. If protein *i* is in *X_U_*, we desire the probability that *z_i_* = *k* for each *k* = 1, …, *K*. That is, for each unlabelled protein, we want the probability of belonging to each component (given a model and the observed data).

We initially model the distribution of profiles associated with proteins that localise to the *k*-th component as multivariate normal with mean vector ***μ**_k_* and covariance matrix Σ_*k*_, so that:

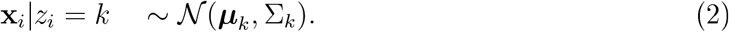

For any *i*, we define the prior probability of the *i*-th protein localising to the *k*-th component to be *p*(*z_i_ = k*) = *π_k_*. Letting 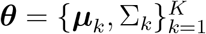 denote the set of all component mean and covariance parameters, and 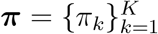 denote the set of all mixture weights, it follows (from the law of total probability) that:

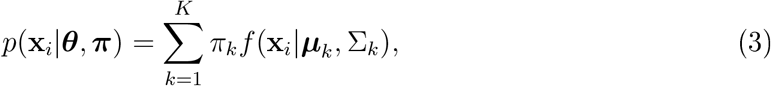

where *f*(**x**|***μ***, Σ) denotes the density of the multivariate normal with mean vector ***μ*** and covariance matrix Σ evaluated at **x**.

Equation (3) defines a generative probabilistic model known as a *mixture model*. Such models are useful for describing populations that are composed of a number of distinct homogeneous subpopulations. In our case, we model the full complement of measured proteins as being composed of *K* subpopulations, each corresponding to a different organelle or sub-cellular compartment. The literature of mixture model applications to biology is rich and some recent example include applications to retroviral integration sites (Kirk *et al.*, 2016), genome-wide associations studies (Liley *et al.*, 2017), single-cell transcriptomics (Lönnberg *et al.*, 2017) and affinity purification MS proteomics (Choi *et al.*, 2010).

Though some proteins are well described as belonging to a single component, many proteins multi-localise or might belong to uncharacterised organelles. In order to allow the model to better account for these “outliers” that cannot be straightforwardly allocated to any single known component, we extend it by introducing an additional “outlier component”. To do this, we augment our model by introducing a further indicator latent variable ø, Each protein **x**_*i*_ is now described by an addition al variable *φ_i_*, with *φ_i_* = 1 indicating that protein **x**_*i*_ belongs to a organelle derived component and *φ_i_* = 0 indicating that protein **x**_*i*_ is not well described by these known components. This outlier component is modelled as a multivariate T distribution with degrees of freedom *κ*, mean vector **M**, and scale matrix *V*, Thus equation (2) becomes

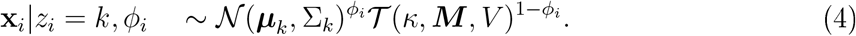

Further let *g*(**x**|*κ*, **M, V**) denote the density of the multivariate T-distribution so that Equation (3) becomes:

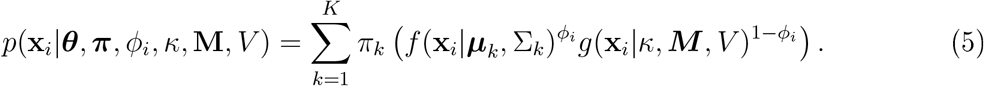

For any *i*, we define the prior probability of the *i*-th protein belonging to the outlier component as *p*(*φ_i_* = 0) = *ϵ*.

We can then rewrite equation (5) in the following way:

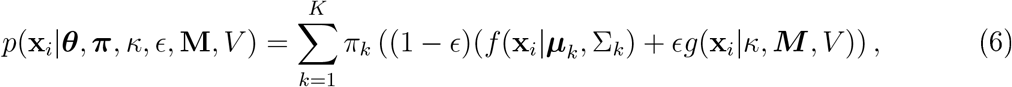

Throughout we take ***κ*** = 4, **M** as the global mean, and *V* as half the global variance of the data, including labelled and unlabelled proteins. The reason for formulating the model as in equation (5) is because it leads to a flexible modelling framework. Furthermore, *φ* has an elegant model selection interpretation, since it decides whether **x**_*i*_ is better modelled bv the known components or the outlier component. It is important to note that *f* and *g* could be replaced by many combinations of distributions and thus could be valuable in modelling other datasets. The choice of parameters for the multivariate T-distribution was decided so that it mimicked a multivariate normal component with the same mean and variance but with heavier tails to better capture dispersed proteins, which we refer to as outlier proteins throughout the text. The variance of the multivariate T-distribution is designed to be large such that is relatively flat when compared with multivariate Gaussian distributions which describe annotated components. Similar approaches for modelling outliers have been explored in the literature and often the outlier term is considered constant or as a Poisson process, independent of the observation (Banfield and Raftery, 1993; Cooke *et al.*, 2011; Coretto and Hennig, 2016; Hennig, 2004).

### 4.2 Model fitting

We adopt a Bayesian approach toward inferring the unknown parameters, 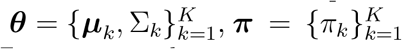, and *ϵ* of the mixture model presented in Equation (5), For ***τ***, we take a conjugate symmetric Dirichlet prior with parameter *β*, so that *τ*_1_, …, *τ_K_* ~ Dirichlet(α); and for the component-specific parameters ***μ**_k_* Σ_*k*_ we take conjugate normal-inverse-Wishart (NIW) priors with parameters {***μ***_0_, λ_0_, *ν*_0_, *S*_0_}, so that:

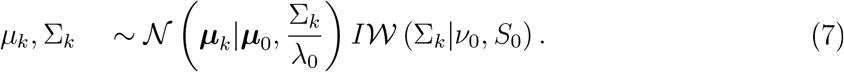

We also place a conjugate Beta prior on e with parameters *u* and *υ*, so that 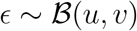. Allowing *ϵ* to be random allows us to infer the number of proteins that are better described by an outlier component rather than any known component.

The full model, which we henceforth refer to as a T-augmented Gaussian Mixture model (TAGM), can then be summarised by the plate diagram shown in Figure 13.

**Figure 13:**
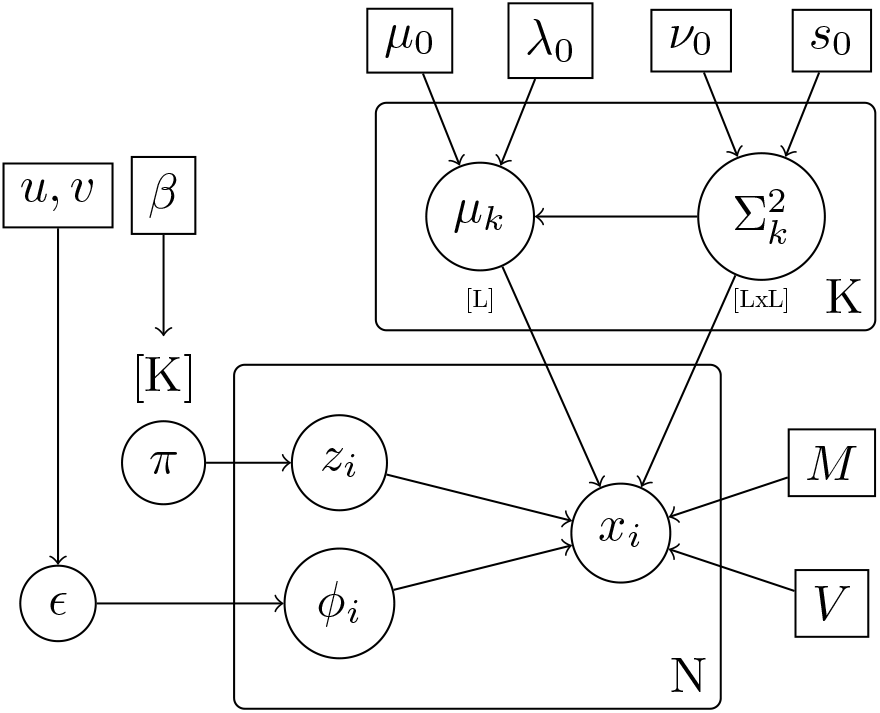
Plate diagram for TAGM model. This diagram specifies the conditional independencies and parameters in our model.

To perform inference for the parameters, we make use of both the labelled and unlabelled data. For the labelled data *X_L_*, sinee *z_i_* and *φ_i_* are known for these proteins, we can update the parameters with their data analytically by exploiting eonjugaey of the priors (see, for example, Gelman *et al.*, 1995), For the unlabelled data we do not have such information and so in the next sections we explain how to make inferences of the latent variables.

### 4.3 Prediction of localisation of unlabelled proteins

Having obtained the posterior distribution of the model parameters analytically using, at first, the labelled data only, we wish to predict the component to which each of the unlabelled proteins belongs. The probability that a protein belongs to any of the *K* known components, that is *z_i_ = k* and *φ_i_* = 1, is given by (see appendix 5.1 for derivations):

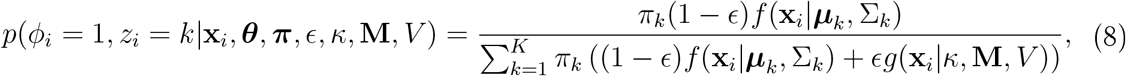

whilst on the other hand,

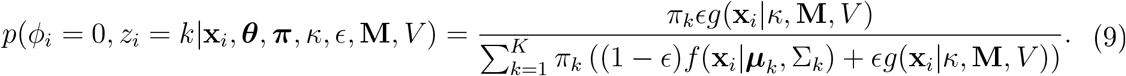

Processing of the unlabelled data can be done by inferring *maximum a posteriori* (MAP) estimates for the parameters. However, this approach fails to account for the uncertainty in the parameters, thus we additionally explore inferring the distribution over these parameters.

#### 4.3.1 Maximum a posteriori prediction

We use the Expectation-Maximisation (EM) algorithm (Dempster *et al.*, 1977) to find *maximum a posteriori* (MAP) estimates for the parameters (see, for example, Murphy, 2012), To specify the parameters of the prior distributions, we use a simple set of heuristics provided by Fraley and Raftery (2007), By defining the following quantities

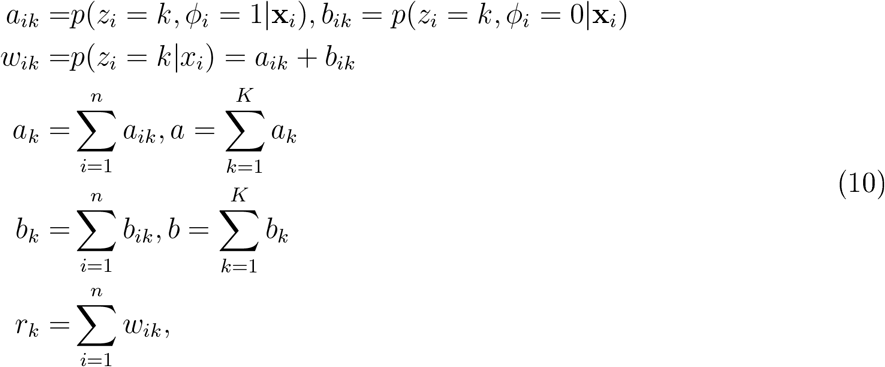

we can compute

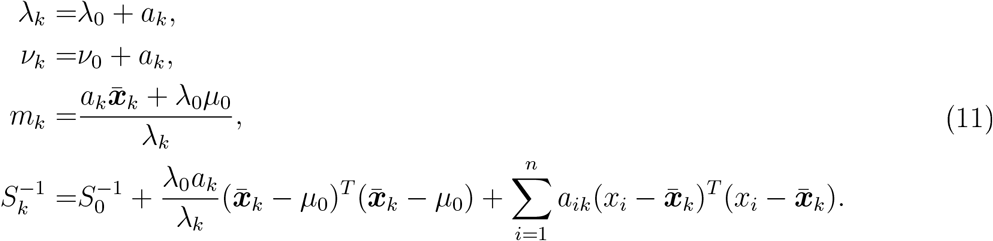

Then the parameters of the posterior mode are:

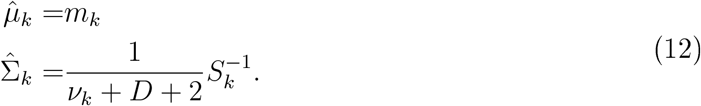

We note if ***x**_i_* is a labelled protein then *a_ik_* = 1 and these parameters can be updated without difficulty. The above equation constitutes a backbone of the E-step of the EM algorithm, with the entire algorithm specified by the following summary:

E-Step: Given the current parameters compute the values given by equations (10), with formulae provided in equations (8) and (9).

M-Step: Compute

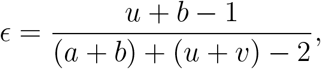

and

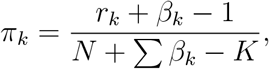

as well as

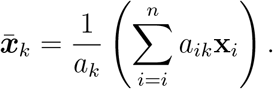

Finally, compute the MAP estimates given by equations (12), These estimates are then used in the following iteration of the E-step.

Denoting by Q the expected value of the log-posterior and letting *t* denote the current iteration of the EM algorithm, we iterate until |*Q*(***θ***|***θ***_*t*_) – *Q*(***θ***|***θ***_*t*−1_)| < *δ*; for some prespecified *δ* > 0. Once we have found MAP estimates for the parameters ***θ**_MAP_*, ***τ**_MAP_* and *ϵ_MAP_* we proceed to perform prediction. We plug the MAP parameter estimates into Equation (8) in order to obtain the posterior probability of protein *i* localising to component *k*, *p*(*z_i_* = *k*,*φ* = 1|**x**_*i*_, ***θ**_map_*, ***τ**_MAP_, ϵ_MAP_,κ*, **M**, *V*), To make a final assignment, we may allocate each protein according to the component that has maximal probability, A full technical derivation of the EM algorithm can be found in the appendix (appendix 5.1).

#### 4.3.2 Uncertainty in the posterior localisation probabilities

The MAP approach described above provides us with a probabilistic assignment, *p*(*z_i_* = *k, φ* = 1|**x**_*i*_, ***θ**_map_*, ***τ**_MAP_, ϵ_MAP_, κ*, **M**, *V*), of each unlabelled protein to each component. However, it fails to account for the uncertainty in the parameters ***θ, τ** and ϵ*. To address this, we can sample parameters from the posterior distribution.

Let 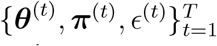 be a set of *T* sampled values for the parameters ***θ, τ** and ϵ*, drawn from the posterior.

The assignment probabilities can then be summarised by the Monte-Carlo average:

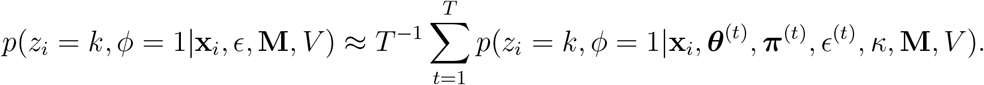

Other summaries of the assignment probabilities can be determined in the usual ways to obtain, for example, interval-estimates. We summarise interval-estimates using the 95% equi-tailed interval, which is defined by the 0.025 and 0.975 quantiles of the distribution of assignment probabilities, 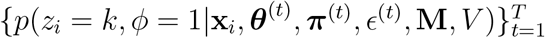.

Sampling parameter values in our model requires us to compute the required conditional probabilities and then a straightforward Gibbs sampler can be used to sample in turn from these conditionals. In addition, we can bypass sampling the parameters by exploiting the conjugacy of our priors. By marginalising parameters in our model we can obtain an efficient collapsed Gibbs sampler and therefore only sample the component allocation probabilities and the outlier allocation probabilities. The derivations and required conditionals can be found in the appendix (appendix 5.2).

### 4.4 Classifier assessment

We compared the classification performance of the two above learning schemes to the K-nearest neighbours (KNN) and the weighted support vector machine (SVM) classifiers.

The following schema was used to assess the classifier performance of all methods. We split the marker sets for each experiment into a class-stratified training (80%) and test (20%) partitions, with the separation formed at random. The true classes of the test profiles are withheld from the classifier, whilst the algorithm is trained. The algorithm is then assessed on its ability to predict the classes of the proteins in the test partition for generalisation accuracy. How each classifier is trained is specific to that classifier. The KNN and SVM have hvperparameters optimised using 5-fold cross-validation. This 80/20 data stratification 100 100 and class specific FI scores (Breckels *et al.*, 2016b). The FI score is the harmonic mean of the precision and recall, more precisely:

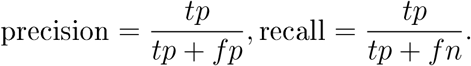

tp denotes the number of true positives; fp the number of false positives and fn the number of false negatives. Thus

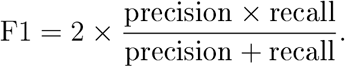

High Macro FI scores indicates that marker proteins in the test dataset are consistently correctly assigned by the classifier. We note that accuracy alone is an inadequate measure of performance, since it fails to quantify false positives.

However, a Bayesian Generative classifier produces probabilistic assignment of observations to classes. Thus while the classifier may make an incorrect assignment it may do so with low probability. The FI score is unforgiving in this situation and will not use this information. To measure this uncertainty, we introduce the quadratic loss which allows us to compare probabilistic assignments (Gneiting and Raftery, 2007). For the SVM, a logistic distribution is fitted using maximum likelihood estimation to the decision values of all binary classifiers. Then, the membership probabilities for the multi-class classification is calculated using quadratic optimisation. The logistic regression model assumes errors which are distributed according to a centred Laplace distribution for the predictions, where maximum likelihood estimation is used to estimate the scale parameter (Meyer *et al.*, 2017). For the KNN classifier, we interpret the proportion of neighbours belonging to each class as a non-parametric posterior probability. To avoid non-zero probabilities for classes we perform Laplace smoothing; that is, the posterior allocation probability is given by

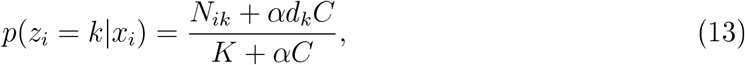

where *N_ik_* is the number of neighbours belonging to class *k* in the neighbourhood of *x_i_ C* is the number of classes, *K* is the number of nearest neighbours (optimised through 5-fold cross validation) and *d_k_* is the incidence rate of each class in the training set. Finally, *α* > 0 is the pseudo-count smoothing parameter. Motivated by a Bayesian interpretation of placing a Jeffrey’s type Dirichlet prior over multinomial counts, we choose *α* = 0.5 (Hazimeh and Zhai, 2015; Valcarce *et al.*, 2016; Manning *et al.*, 2008). The quadratic loss is given by the following formula:

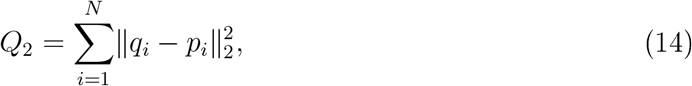

where ‖·‖_2_ is the *l*_2_ norm and *q_i_* is the true classification vector and *pi* is a vector of predicted assignments to each class. It is useful to note that the corresponding risk function is the mean square error (MSE), which is the expected value of the quadratic loss.

## Acknowledgments

The authors would also like to thank Dr Sean B. Holden, University of Cambridge, for helpful discussions.

## 5 Appendices

### 5.1 Appendix 1: Derivation of EM algorithm for TAGM model

This appendix give a formal derivation of the EM algorithm used for our model. Computations are standard but useful and similar technical summaries can be found (for example see Fraley and Raftery (2005); Murphy (2007)) We let *H* = {***μ***_0_, λ_0_, *ν*_0_, *S*_0_} denote the parameters of the normal-inverse-Wishart prior. More precisely:

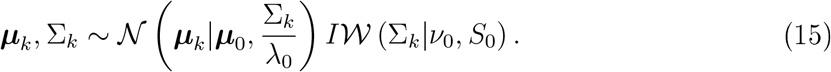

Furthermore, let ***θ_k_*** = {***μ**_k_*, Σ_*k*_}, and let Θ = {*κ*, **M**, *V*} be the parameters of the global 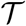 distribution. We specify the following hierarchical Bayesian model.

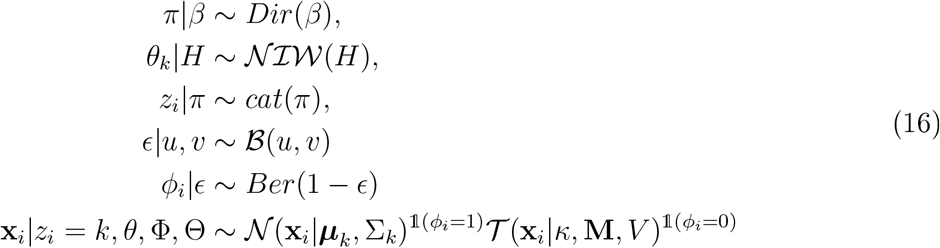

Since *p*(*φ_i_* = 1) = 1 − *ϵ*, we can rewrite the last line of the model (16) as the following:

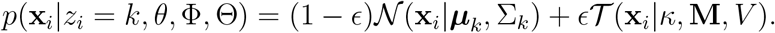

The total joint probability is

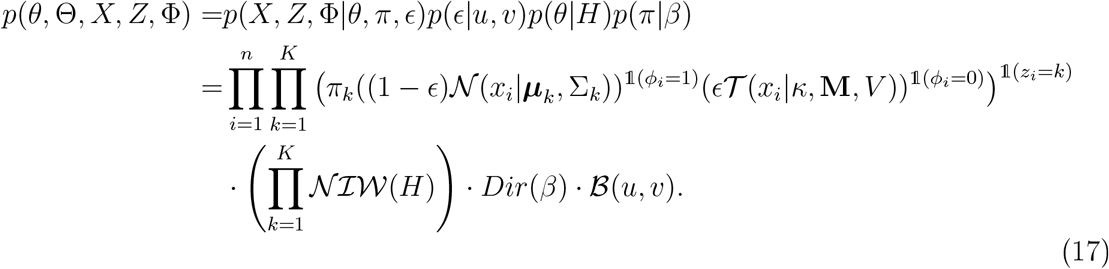

Before we formally derive an EM algorithm for this model, we derive a few useful quantities, Let *f*(**x|*μ***, Σ) denote the density of the multivariate normal with mean vector ***μ*** and covariance matrix Σ evaluated at **x** and further let *g*(**x**|*κ*, **M**, *V*) denote the density of the multivariate T-distribution, We compute that

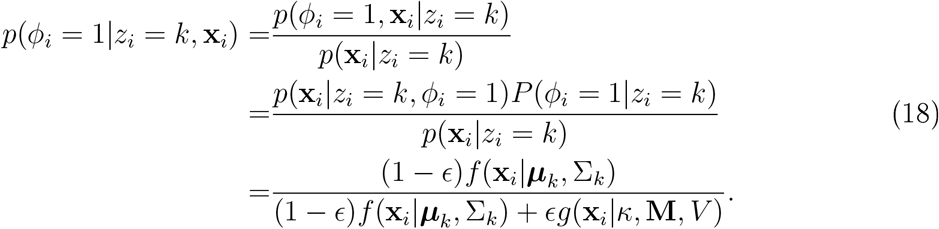

Likewise we see that,

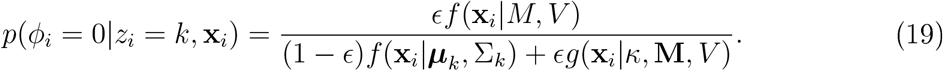

Thus

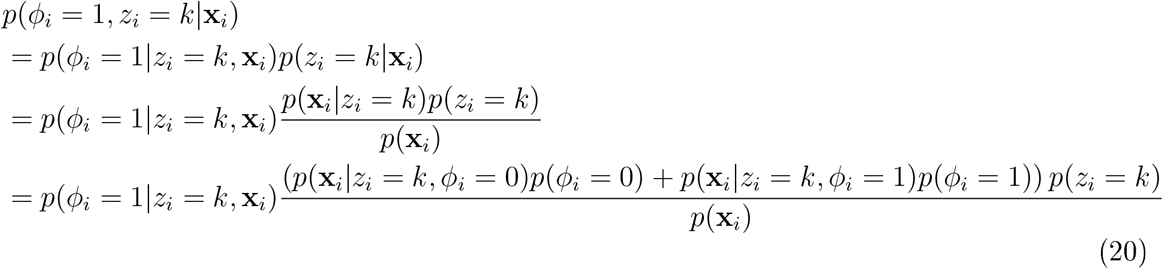

and then substituting values leads to

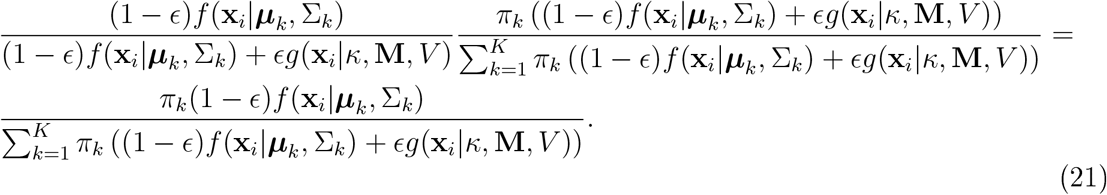

We also see that

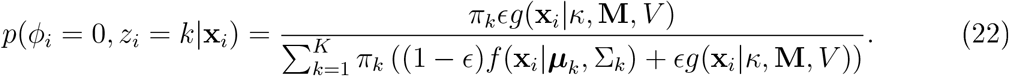

We can now formally derive the EM algorithm for this model. First, we compute the expected value of the log-posterior function with respect to the conditional distribution of the latent variable given the observations (under the current estimate of the parameters). For notational convenience we suppress the dependence on the parameters.

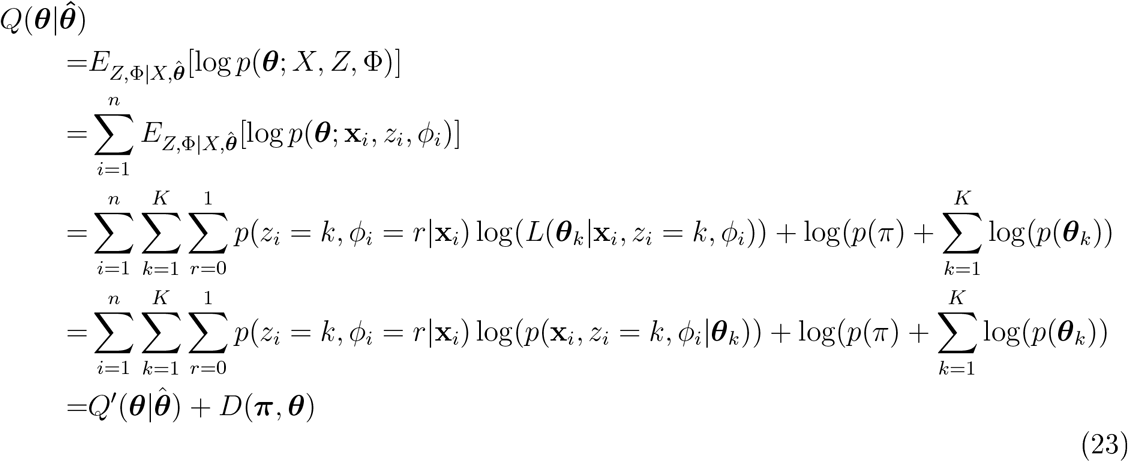

We note that the equation splits up into a likelihood term *Q′* plus the log prior *D*, The coefficient of the first term in the equation above has already been derived and the other term is given by:

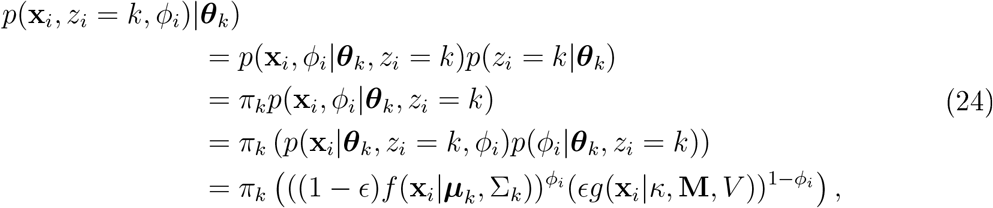

where we used that *φ_i_* was a binary random variable. Thus we see that

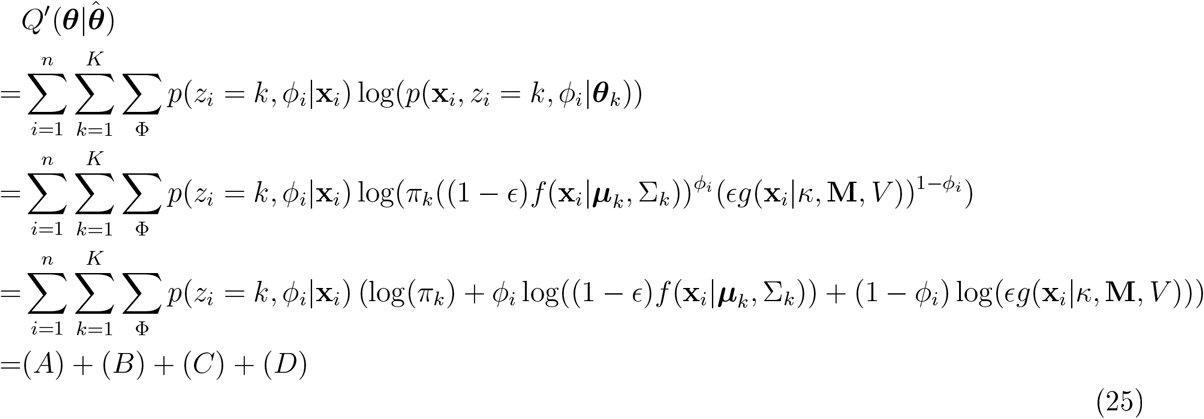

where

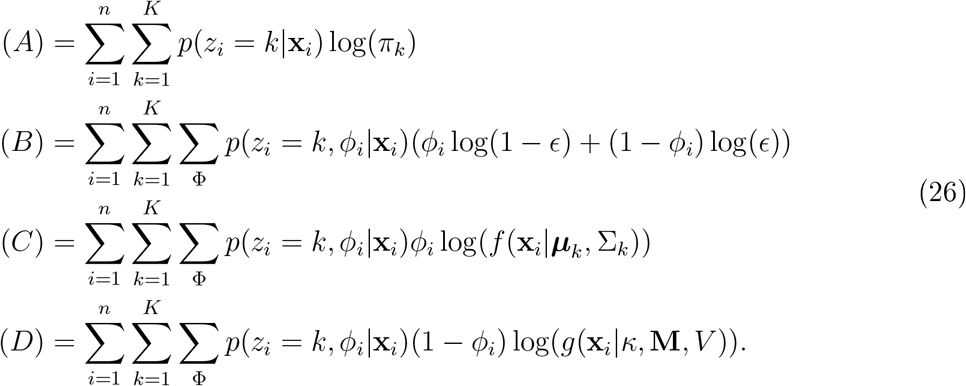

Then again using that *φ_i_* is binary we can make the following simplifications.

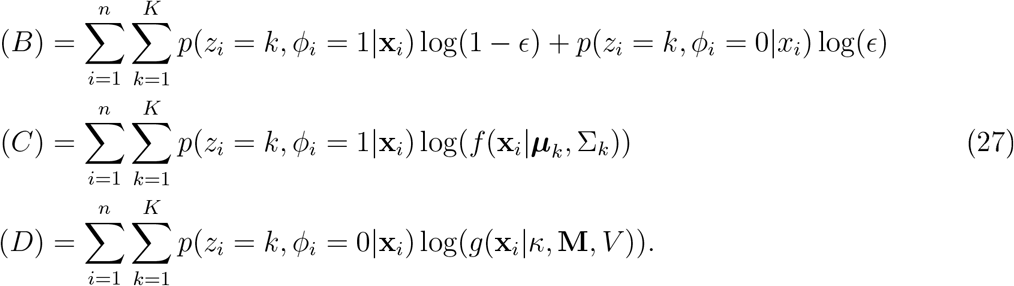

Terms can now be maximised by considering terms independently because of linearity, Note that the equations 8 and 9 are computed with respect to the current estimated values of the parameters. For convenience set the following notation

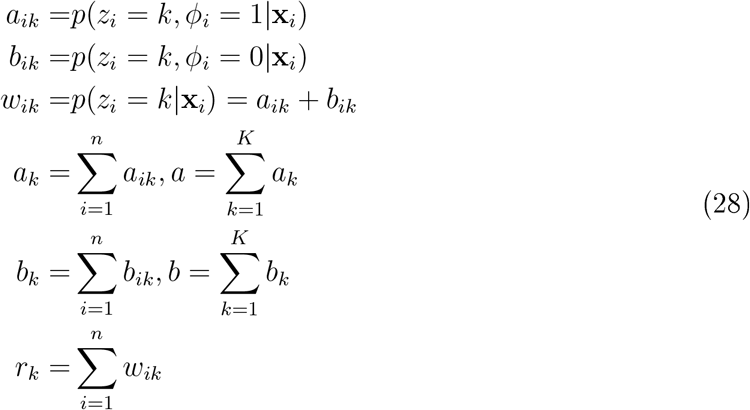

The maximisation step requires finding 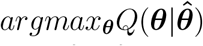, this can be found for parameter separately for each linear term. To find 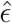, we need only consider computing the maximisation step from equation (B), First set *ϵ*_1_ = 1 − *ϵ* and *ϵ*_2_ = *ϵ* and add the log prior term to equation (B), Thus, the required Lagrangian is

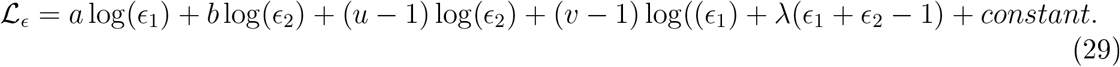

Solving this system leads to

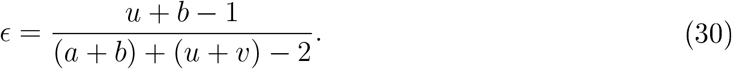

To find the MAP estimate for ***τ***, we examine equation (A) and add the log prior. Furthermore we must maximise ***τ*** under the constraint that 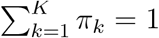 The Lagrangian for this constrained optimisation problem is the following,

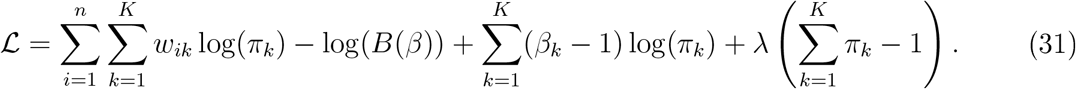

The fixed point of this Lagrangian solves the required constrained optimisation problem and *B*(*β*) denotes the Beta function with parameter *α*.

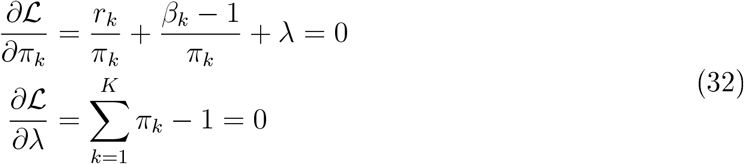

Solving this pair of equations yields

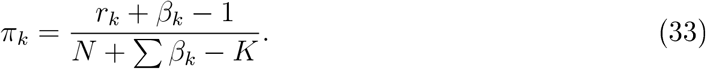

To find the posterior mode of the remaining parameters requires some work. First we recall that the normal inverse-Wishart prior is proportional to:

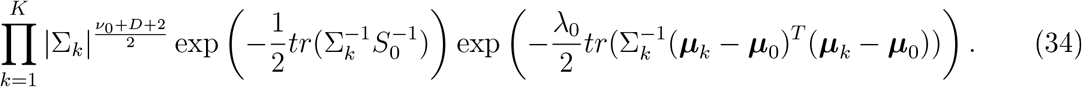

The required equation we are interested in is (C).

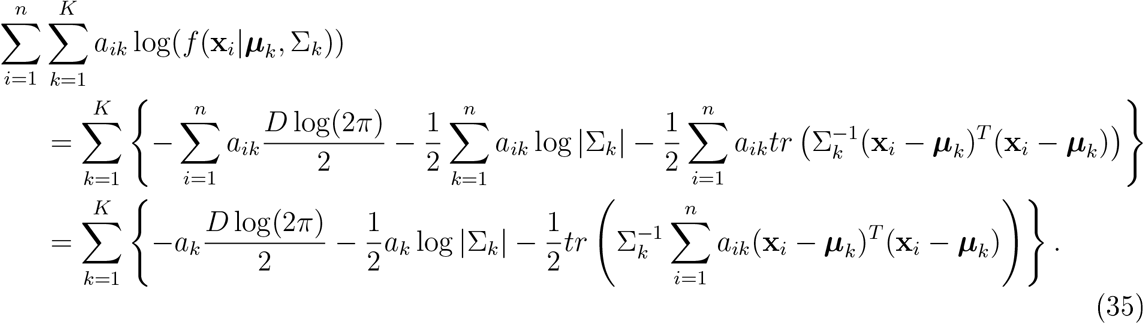

Now to derive the M-step objective we remove the constant terms and add on the log prior. This leads to

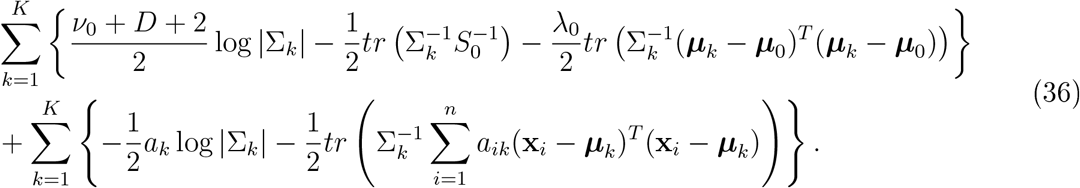

This can be rewritten as

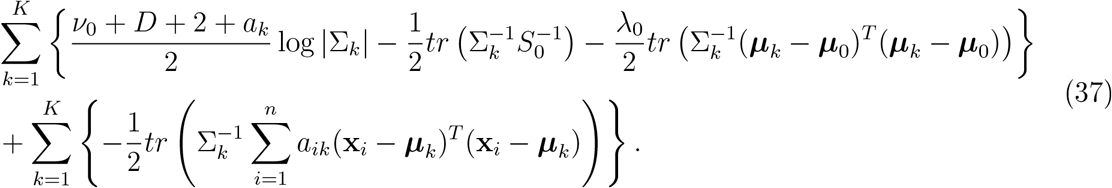

Now define 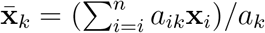 and note the following algebraic rearrangements.

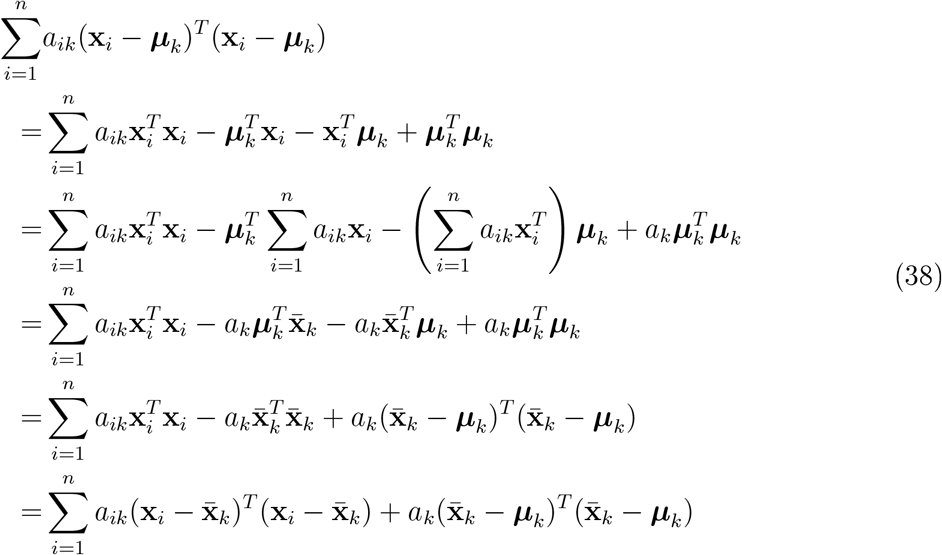

This allows us to rewrite equation 37 as

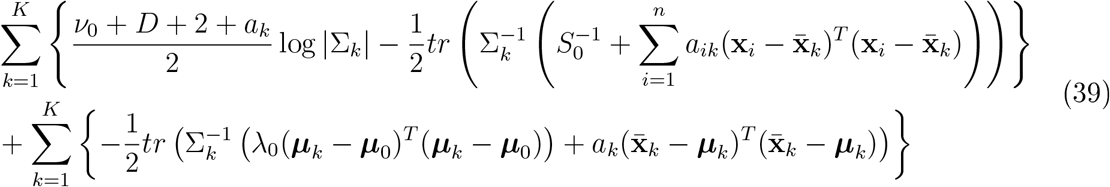

This can be written as:

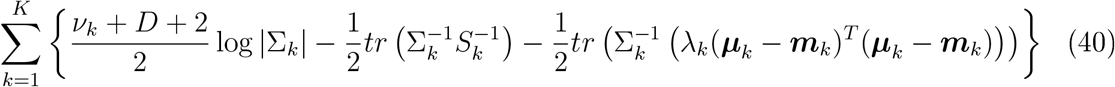

where,

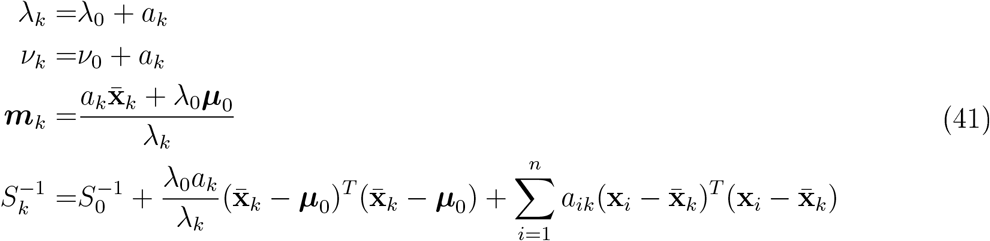

Thus the parameters of the posterior mode are:

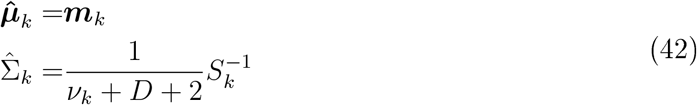

To summarise the EM algorithm, we iterate between the two steps:

E-Step: Given the current parameters compute the values given by equations (28), with formulas provided in equations (8) and (9).

M-Step: Compute

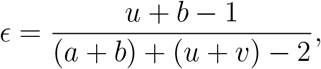

and

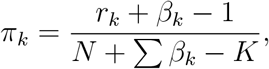

as well as

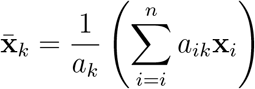

Compute the MAP estimates given by equations (42), These estimates are then used in the following iteration of the E-step, Iterate until |Q(6|6_t_) — Q(6|6_t-1_)| < ó for some pre-specified ó > 0,

### 5.2 Appendix 2: Derivation of collapsed Gibbs sampler for TAGM model

To derive the Gibbs sampler we write down all the conditional probabilities. Then, exploiting conjugacy, we can marginalise parameters in the model. Recall the total joint probability is the following:

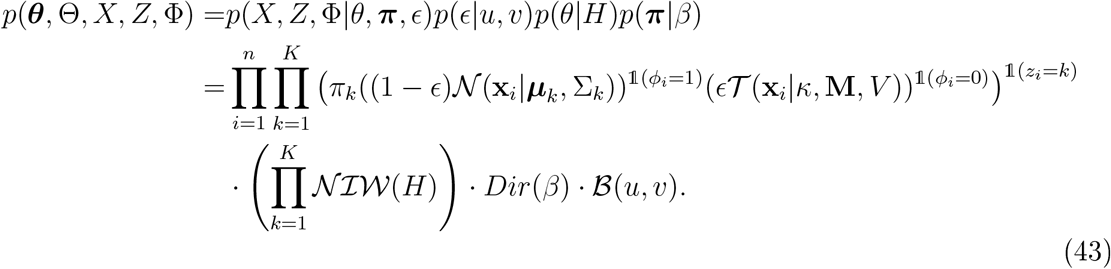

Suppose we know the hidden latent component allocations *z_i_* and outlier allocations *φ_i_*. Then we could sample from the a required normal distribution. The conditional probability of the parameters given the allocations is given by:

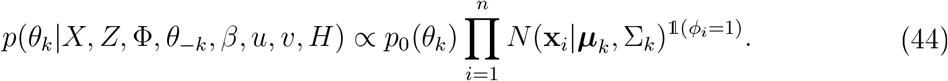

The prior is conjugate and so the posterior belongs to the same parametric family as the prior, a NIW distribution, and so the parameters can be updated as follows:

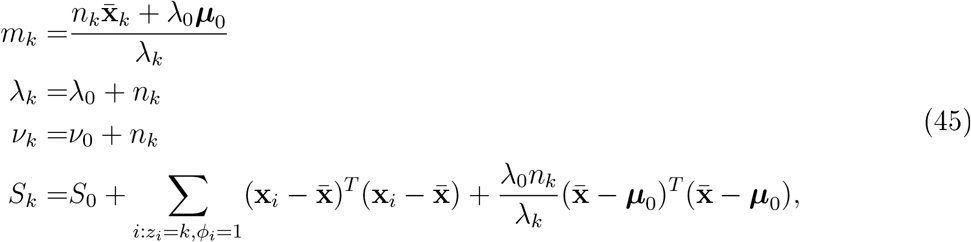

where *n_k_* = |{**x**_*i*_|*z_i_* = *k*, *φ_i_* = 1}|. Now we write down the conditional of the component allocations

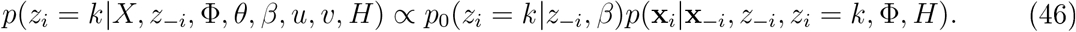

The first term in this equation is

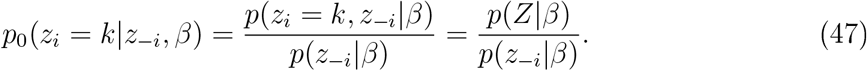

To calculate the numerator we proceed by marginalising over ***τ*** as follows

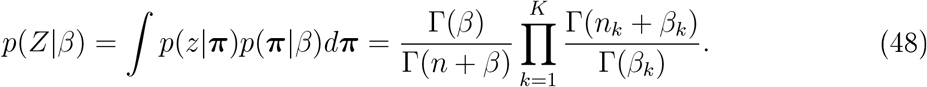

Hence, we arrive at the following probability:

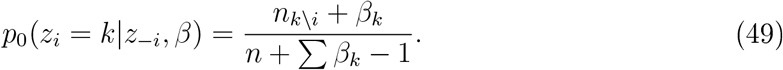

The conditional for the second term of 46 is more tricky. First note the following conditional distributions

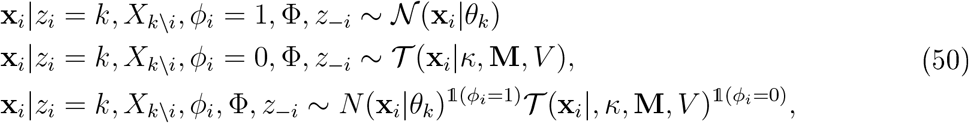

where we denote *X_k\i_* as the observations associated with class *k*, besides *x_i_*. Now, we first note that:

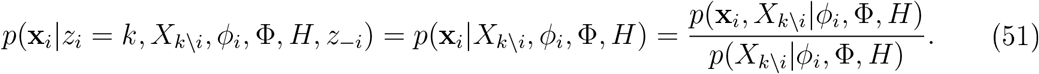

Thus, we find an equation for the numerator, using the fact that terms associated with *φ_i_* = 0 do not depend on *k* and thus can be absorbed into the normalising constant.

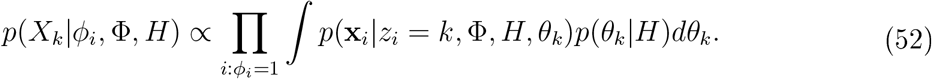

This is the marginal likelihood of the data. Thus the ratio in 51 is the posterior predictive which is given by the non-centred T-distribution with formula given by:

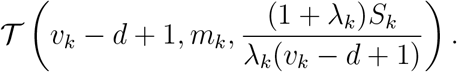

Thus, we can compute the following:

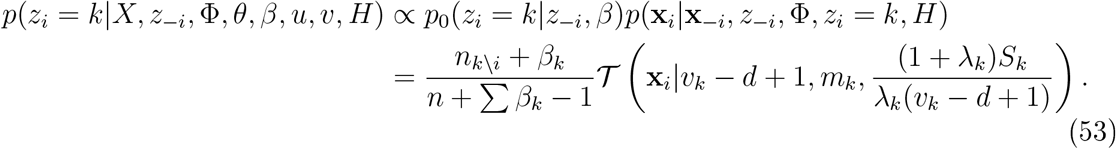

It remains to compute the conditional for the *φ_i_*. By first recalling that *φ_i_* is binary we see that

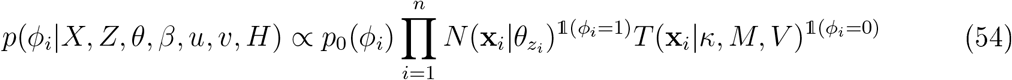

can be written as

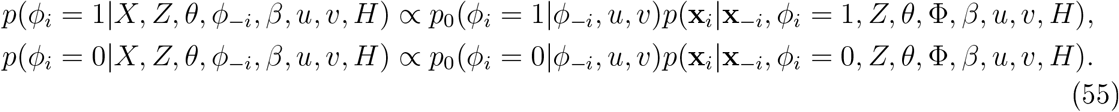

First we need to compute a formula for *p*_0_(*φ_i_*|*φ_−i_*, *u, υ*), First we see that

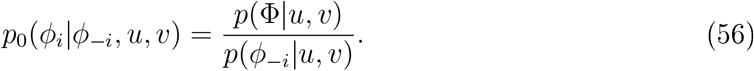

The numerator can be computed bv marginalising over *ϵ*:

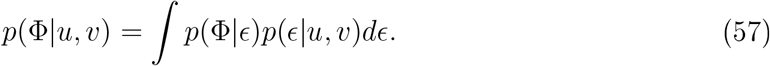

We denote 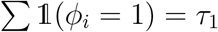 and 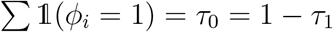. Then it is easy to see that

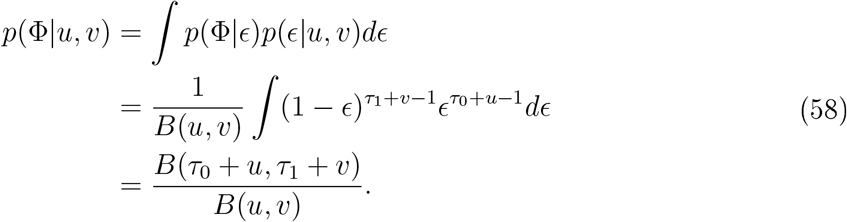

Hence,

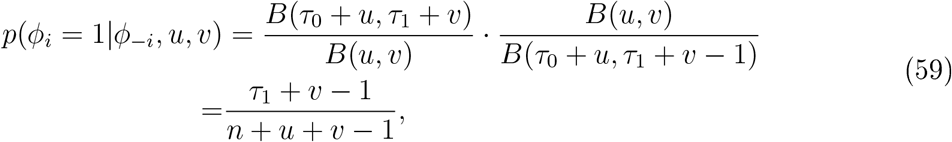

where *n* = *τ*_1_ + *τ*_2_. In general,

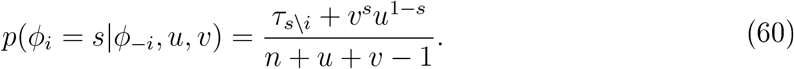

Now we return to computing *p*(**x**_*i*_|**x**_−*i*_, *Z, θ, φ_i_* = 1, Φ, *α, u, υ, H*), First we see that

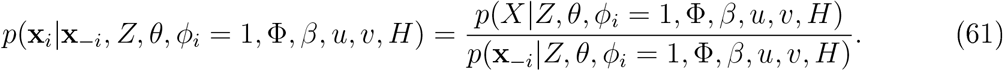

Thus if we integrate over the parameters, we would have a ratio of marginal likelihoods giving the posterior predictive which is a non-centred T-distribution:

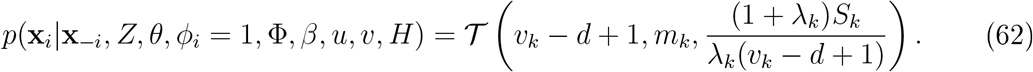

In the other case that *φ* = 0, we have that

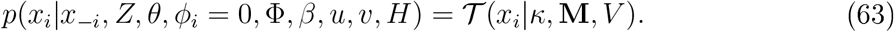

Thus we can compute:

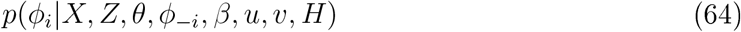

and sample from the required distribution. Thus, we can summarise the collapsed Gibbs sampler as follows:

1. Update the priors with the labelled data
2. For the unlabelled observations, in turn, compute the probability of assigning to each component
3. Sample a label according to this probability
4. Compute the probability of belonging to this class or the outlier component
5. Sample an indicator to a class specific component or the outlier component
6. If we assign to the class specific component update the class specific posterior distribution with the statistics of this observation
7. Update other posteriors as appropriate.
8. Once all unlabelled observations have a been assigned, consider the observations sequentially, removing the statistics from the posteriors and then performing steps 2-7, We repeat this process for all unlabelled observations.
9. repeat 7-8 until convergence of the Markov-ehain.

The computational bottleneck in the algorithm is computing the posterior updates for the parameters

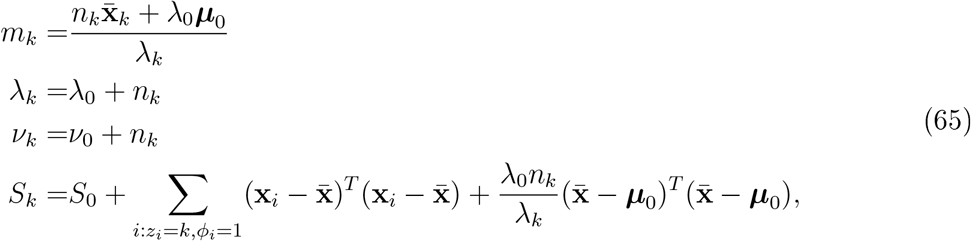

We first note that

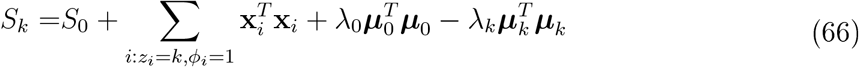

Let us denote 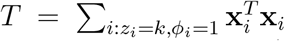. Thus we can derive a set of iterative updates to speed up computation when adding/removing statistics from clusters. More precisely, indicating updated posterior parameters by a prime, if we remove statistics of observation *i* from cluster *k*, we see that

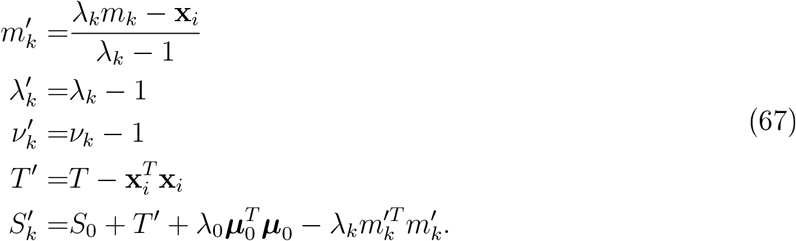

Likewise if we add the statistics of observation *i* to cluster *k*, we see that

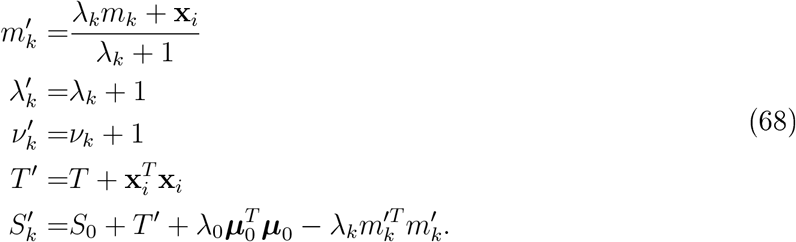

### 5.3 Appendix 3: Convergence diagnostics of EM algorithm

**Figure 14:**
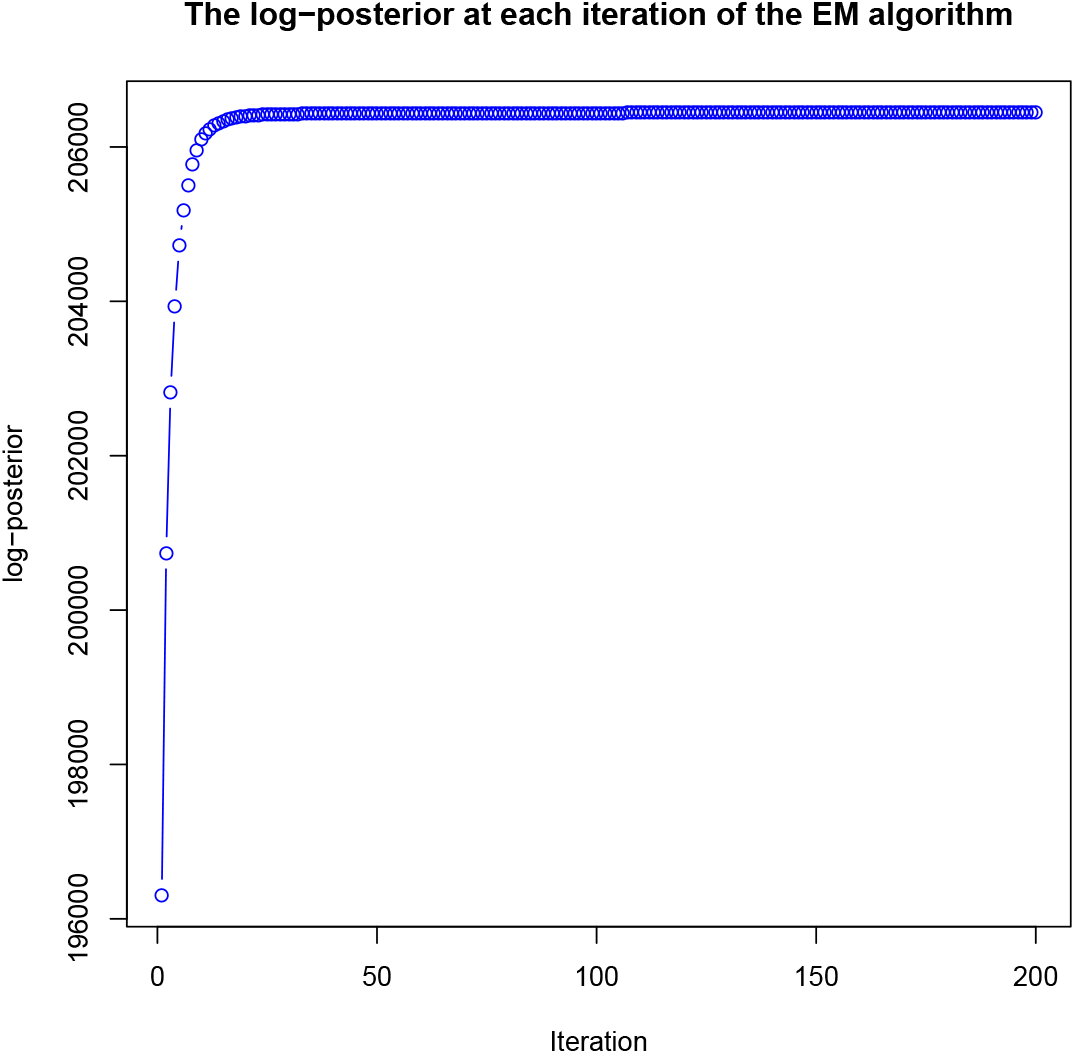
Plot of the log-posterior at each iteration of the EM algorithm to demonstrate monotonirìty and convergence

### 5.4 Appendix 4: Trace plots for assessing MCMC convergence

**Figure 15:**
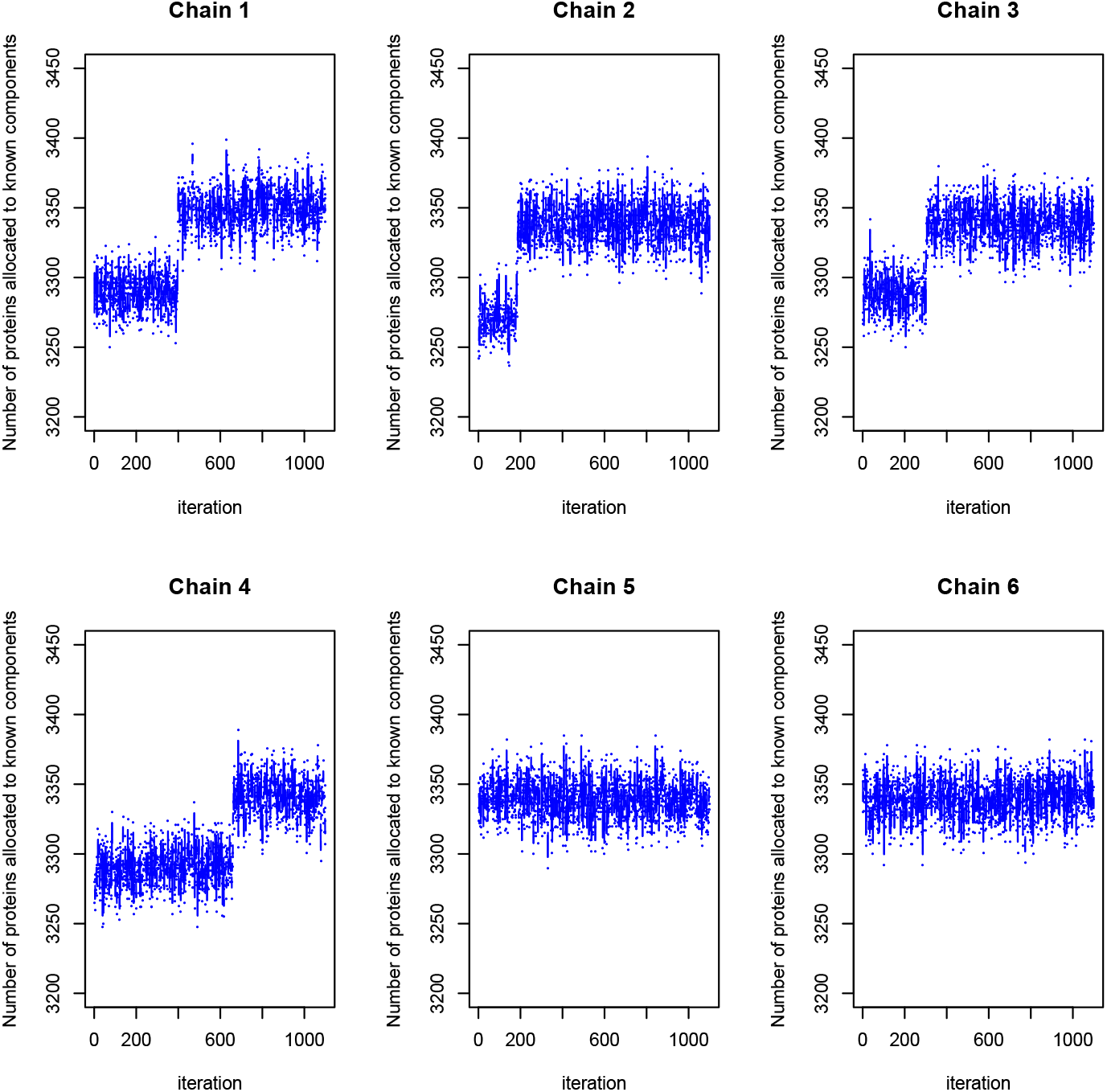
Trace plots of the number of proteins allocated to the known components in each of 6 parallel MCMC runs. Chain 4 is discarded because of lack of convergence. 600 samples are retained from remaining chains and pooled.

### 5.5 Appendix 5: F1 t-tests

**Table 2:** Adjusted P-values for pairwise T-tests for Macro F-1 score classifier evaluation on the Drosophila dataset

|  | SVM | KNN | MAP |
| --- | --- | --- | --- |
| KNN | 2.7E-03 |  |  |
| MAP | 3.3E-02 | 3.4E-01 |  |
| MCMC | 3.4E-01 | 3.3E-02 | 2.3E-01 |

**Table 3:**
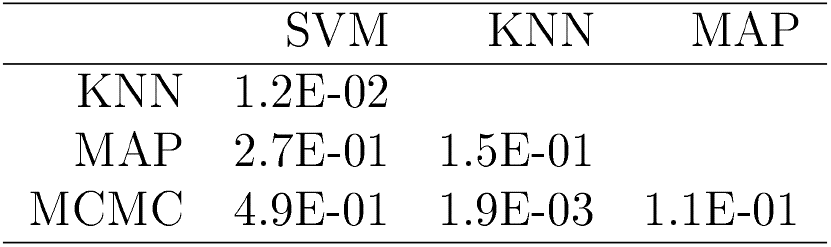
Adjusted P-values for pairwise T-tests for Macro F-1 score classifier evaluation on the Chicken DT40 dataset

|  | SVM | KNN | MAP |
| --- | --- | --- | --- |
| KNN | 1.2E-02 |  |  |
| MAP | 2.7E-01 | 1.5E-01 |  |
| MCMC | 4.9E-01 | 1.9E-03 | 1.1E-01 |

**Table 4:**
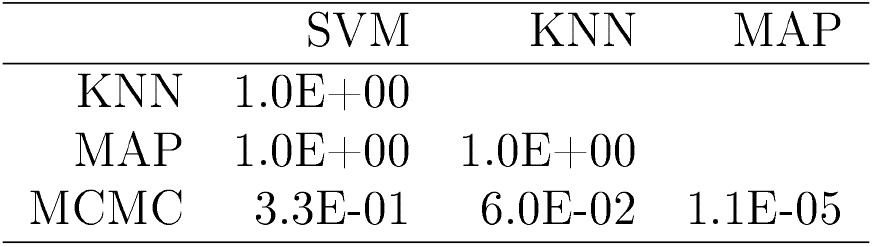
Adjusted P-values for pairwise T-tests for Macro F-1 score classifier evaluation on the mouse dataset

**Table 5:** Adjusted P-values for pairwise T-tests for Macro F-1 score classifier evaluation on the HeLa dataset

|  | SVM | KNN | MAP |
| --- | --- | --- | --- |
| KNN | 1.4E-35 |  |  |
| MAP | 3.3E-06 | 6.7E-21 |  |
| MCMC | 8.0E-59 | 3.2E-91 | 2.4E-70 |

**Table 6:** Adjusted P-values for pairwise T-tests for Macro F-1 score classifier evaluation on the U2-OS dataset

|  | SVM | KNN | MAP |
| --- | --- | --- | --- |
| KNN | 1.3E-02 |  |  |
| MAP | 4.3E-04 | 3.3E-09 |  |
| MCMC | 5.8E-01 | 3.5E-03 | 3.1E-03 |

**Table 7:** Adjusted P-values for pairwise T-tests for Macro F-1 score classifier evaluation on the HeLa wild (Hirst et al.) dataset

|  | SVM | KNN | MAP |
| --- | --- | --- | --- |
| KNN | 2.2E-08 |  |  |
| MAP | 1.0E-34 | 6.8E-14 |  |
| MCMC | 7.4E-05 | 5.3E-02 | 1.0E-20 |

**Table 8:** Adjusted P-values for pairwise T-tests for Macro F-1 score classifier evaluation on the HeLa KOI (Hirst et al.) dataset

|  | SVM | KNN | MAP |
| --- | --- | --- | --- |
| KNN | 5.3E-02 |  |  |
| MAP | 1.7E-23 | 7.9E-27 |  |
| MCMC | 9.1E-02 | 5.8E-04 | 1.8E-19 |

**Table 9:** Adjusted P-values for pairwise T-tests for Macro F-1 score classifier evaluation on the HeLa KO2 (Hirst et al.) dataset

|  | SVM | KNN | MAP |
| --- | --- | --- | --- |
| KNN | 1.3E-01 |  |  |
| MAP | 1.1E-55 | 1.1E-55 |  |
| MCMC | 1.0E-18 | 6.3E-22 | 2.0E-26 |

**Table 10:** Adjusted P-values for pairwise T-tests for Macro F-1 score classifier evaluation on the Primary Fibroblasts Mock 24hpi dataset

|  | SVM | KNN | MAP |
| --- | --- | --- | --- |
| KNN | 9.6E-02 |  |  |
| MAP | 4.1E-07 | 1.1E-09 |  |
| MCMC | 2.8E-27 | 1.0E-28 | 6.3E-10 |

**Table 11:** Adjusted P-values for pairwise T-tests for Macro F-1 score classifier evaluation on the Primary Fibroblasts Mock 48hpi dataset

|  | SVM | KNN | MAP |
| --- | --- | --- | --- |
| KNN | 6.6E-07 |  |  |
| MAP | 1.3E-10 | 2.0E-01 |  |
| MCMC | 1.6E-05 | 2.0E-01 | 6.2E-03 |

**Table 12:** Adjusted P-values for pairwise T-tests for Macro F-1 score classifier evaluation on the Primary Fibroblasts Mock 72hpi dataset

|  | SVM | KNN | MAP |
| --- | --- | --- | --- |
| KNN | 3.9E-03 |  |  |
| MAP | 9.5E-01 | 8.6E-03 |  |
| MCMC | 6.4E-02 | 3.0E-01 | 8.6E-02 |

**Table 13:** Adjusted P-values for pairwise T-tests for Macro F-1 score classifier evaluation on the Primary Fibroblasts Mock 96hpi dataset

|  | SVM | KNN | MAP |
| --- | --- | --- | --- |
| KNN | 8.6E-03 |  |  |
| MAP | 1.1E-02 | 8.6E-01 |  |
| MCMC | 3.7E-06 | 1.6E-02 | 3.3E-02 |

**Table 14:** Adjusted P-values for pairwise T-tests for Macro F-1 score classifier evaluation on the Primary Fibroblasts Mock 120hpi dataset

|  | SVM | KNN | MAP |
| --- | --- | --- | --- |
| KNN | 1.9E-23 |  |  |
| MAP | 1.4E-02 | 2.3E-34 |  |
| MCMC | 3.8E-07 | 1.6E-81 | 2.0E-02 |

**Table 15:** Adjusted P-values for pairwise T-tests for Macro F-1 score classifier evaluation on the Primary Fibroblasts HCMV 24hpi dataset

|  | SVM | KNN | MAP |
| --- | --- | --- | --- |
| KNN | 4.6E-01 |  |  |
| MAP | 2.6E-05 | 1.7E-04 |  |
| MCMC | 1.7E-04 | 1.3E-03 | 5.5E-01 |

**Table 16:** Adjusted P-values for pairwise T-tests for Macro F-1 score classifier evaluation on the Primary Fibroblasts HCMV 48hpi dataset

|  | SVM | KNN | MAP |
| --- | --- | --- | --- |
| KNN | 1.0E-02 |  |  |
| MAP | 4.6E-01 | 1.5E-03 |  |
| MCMC | 1.2E-02 | 7.3E-01 | 1.5E-03 |

**Table 17:** Adjusted P-values for pairwise T-tests for Macro F-1 score classifier evaluation on the Primary Fibroblasts HCMV 72hpi dataset

|  | SVM | KNN | MAP |
| --- | --- | --- | --- |
| KNN | 5.5E-02 |  |  |
| MAP | 9.5E-06 | 3.4E-02 |  |
| MCMC | 1.1E-01 | 6.2E-01 | 6.4E-03 |

**Table 18:** Adjusted P-values for pairwise T-tests for Macro F-1 score classifier evaluation on the Primary Fibroblasts HCMV 96hpi dataset

|  | SVM | KNN | MAP |
| --- | --- | --- | --- |
| KNN | 2.8E-01 |  |  |
| MAP | 2.6E-09 | 7.2E-08 |  |
| MCMC | 4.2E-10 | 5.6E-09 | 5.7E-01 |

**Table 19:** Adjusted P-values for pairwise T-tests for Macro F-1 score classifier evaluation on the Primary Fibroblasts HCMV 120hpi dataset

|  | SVM | KNN | MAP |
| --- | --- | --- | --- |
| KNN | 2.3E-04 |  |  |
| MAP | 7.1E-04 | 3.8E-10 |  |
| MCMC | 1.4E-01 | 5.7E-02 | 6.0E-05 |

**Table 20:** Adjusted P-values for pairwise T-tests for Macro F-1 score classifier evaluation on the E14TG2a dataset

|  | SVM | KNN | MAP |
| --- | --- | --- | --- |
| KNN | 6.7E-06 |  |  |
| MAP | 6.3E-05 | 4.4E-01 |  |
| MCMC | 4.4E-01 | 6.7E-06 | 8.3E-05 |

### 5.6 Appendix 6: Quadratic loss t-tests

**Table 21:**
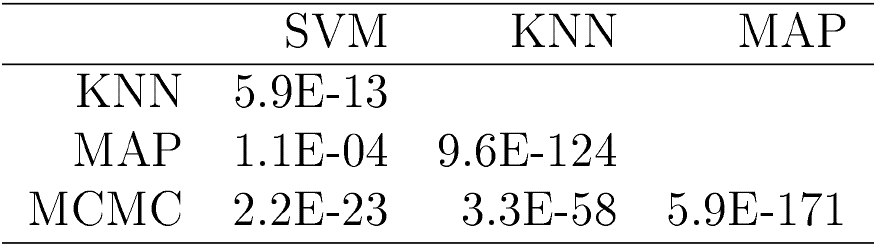
Adjusted P-values for pairwise T-tests for Quadratic Loss classifier evaluation on the Drosphila dataset

|  | SVM | KNN | MAP |
| --- | --- | --- | --- |
| KNN | 5.9E-13 |  |  |
| MAP | 1.1E-04 | 9.6E-124 |  |
| MCMC | 2.2E-23 | 3.3E-58 | 5.9E-171 |

**Table 22:**
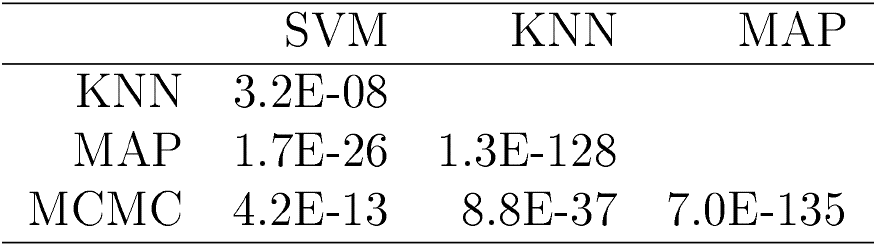
Adjusted P-values for pairwise T-tests for Quadratic Loss classifier evaluation on the Chicken DT40 dataset

|  | SVM | KNN | MAP |
| --- | --- | --- | --- |
| KNN | 3.2E-08 |  |  |
| MAP | 1.7E-26 | 1.3E-128 |  |
| MCMC | 4.2E-13 | 8.8E-37 | 7.0E-135 |

**Table 23:** Adjusted P-values for pairwise T-tests for Quadratic Loss classifier evaluation on the mouse dataset

|  | SVM | KNN | MAP |
| --- | --- | --- | --- |
| KNN | 5.5E-14 |  |  |
| MAP | 3.0E-25 | 6.3E-128 |  |
| MCMC | 7.4E-26 | 1.7E-129 | 1.6E-14 |

**Table 24:** Adjusted P-values for pairwise T-tests for Quadratic Loss classifier evaluation on the HeLa dataset

|  | SVM | KNN | MAP |
| --- | --- | --- | --- |
| KNN | 1.2E-02 |  |  |
| MAP | 9.4E-07 | 7.4E-86 |  |
| MCMC | 5.5E-08 | 2.7E-89 | 2.4E-12 |

**Table 25:** Adjusted P-values for pairwise T-tests for Quadratic Loss classifier evaluation on the U2-OS dataset

|  | SVM | KNN | MAP |
| --- | --- | --- | --- |
| KNN | 6.8E-02 |  |  |
| MAP | 7.4E-17 | 1.1E-73 |  |
| MCMC | 1.4E-20 | 6.7E-81 | 8.3E-41 |

**Table 26:** Adjusted P-values for pairwise T-tests for Quadratic Loss classifier evaluation on the HeLa wild (Hirst et al.) dataset

|  | SVM | KNN | MAP |
| --- | --- | --- | --- |
| KNN | 2.3E-92 |  |  |
| MAP | 9.0E-13 | 2.4E-83 |  |
| MCMC | 6.6E-19 | 3.0E-81 | 1.1E-01 |

**Table 27:** Adjusted P-values for pairwise T-tests for Quadratic Loss classifier evaluation on the HeLa KO1 (Hirst et al.) dataset

|  | SVM | KNN | MAP |
| --- | --- | --- | --- |
| KNN | 5.2E-97 |  |  |
| MAP | 1.4E-02 | 1.2E-90 |  |
| MCMC | 2.3E-09 | 7.0E-95 | 2.2E-02 |

**Table 28:** Adjusted P-values for pairwise T-tests for Quadratic Loss classifier evaluation on the HeLa KO2 (Hirst et al.) dataset

|  | SVM | KNN | MAP |
| --- | --- | --- | --- |
| KNN | 8.9E-93 |  |  |
| MAP | 3.1E-01 | 8.1E-91 |  |
| MCMC | 9.0E-06 | 1.5E-83 | 8.9E-05 |

**Table 29:** Adjusted P-values for pairwise T-tests for Quadratic Loss classifier evaluation on the Primary Fibroblasts Mock 24hpi dataset

|  | SVM | KNN | MAP |
| --- | --- | --- | --- |
| KNN | 6.1E-13 |  |  |
| MAP | 1.4E-18 | 4.4E-81 |  |
| MCMC | 3.2E-18 | 7.2E-77 | 5.9E-03 |

**Table 30:** Adjusted P-values for pairwise T-tests for Quadratic Loss classifier evaluation on the Primary Fibroblasts Mock 48hpi dataset

|  | SVM | KNN | MAP |
| --- | --- | --- | --- |
| KNN | 6.1E-18 |  |  |
| MAP | 3.6E-24 | 2.2E-57 |  |
| MCMC | 1.4E-24 | 3.6E-61 | 3.6E-04 |

**Table 31:** Adjusted P-values for pairwise T-tests for Quadratic Loss classifier evaluation on the Primary Fibroblasts Mock 72hpi dataset

|  | SVM | KNN | MAP |
| --- | --- | --- | --- |
| KNN | 1.2E-15 |  |  |
| MAP | 4.5E-23 | 2.5E-89 |  |
| MCMC | 4.2E-23 | 5.1E-91 | 4.4E-01 |

**Table 32:** Adjusted P-values for pairwise T-tests for Quadratic Loss classifier evaluation on the Primary Fibroblasts Mock 96hpi dataset

|  | SVM | KNN | MAP |
| --- | --- | --- | --- |
| KNN | 1.8E-13 |  |  |
| MAP | 1.4E-20 | 3.6E-126 |  |
| MCMC | 5.0E-20 | 1.5E-109 | 5.3E-07 |

**Table 33:** Adjusted P-values for pairwise T-tests for Quadratic Loss classifier evaluation on the Primary Fibroblasts Mock 120hpi dataset

|  | SVM | KNN | MAP |
| --- | --- | --- | --- |
| KNN | 6.7E-14 |  |  |
| MAP | 1.0E-19 | 2.6E-45 |  |
| MCMC | 8.0E-20 | 2.4E-45 | 2.5E-02 |

**Table 34:** Adjusted P-values for pairwise T-tests for Quadratic Loss classifier evaluation on the Primary Fibroblasts HCMV 24hpi dataset

|  | SVM | KNN | MAP |
| --- | --- | --- | --- |
| KNN | 6.0E-22 |  |  |
| MAP | 2.8E-27 | 6.4E-53 |  |
| MCMC | 1.4E-27 | 1.5E-56 | 3.0E-03 |

**Table 35:**
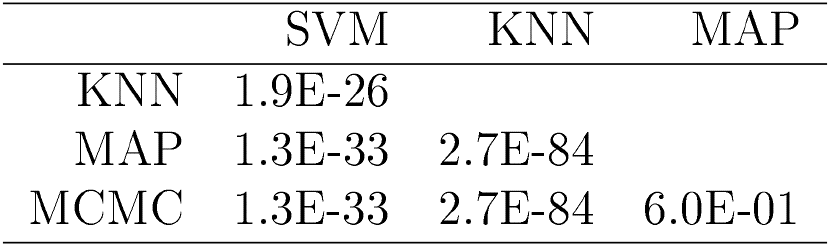
Adjusted P-values for pairwise T-tests for Quadratic Loss classifier evaluation on the Primary Fibroblasts HCMV 48hpi dataset

**Table 36:** Adjusted P-values for pairwise T-tests for Quadratic Loss classifier evaluation on
the Primary Fibroblasts HCMV 72hpi dataset

|  | SVM | KNN | MAP |
| --- | --- | --- | --- |
| KNN | 6.3E-20 |  |  |
| MAP | 1.9E-25 | 2.7E-57 |  |
| MCMC | 1.2E-25 | 3.4E-58 | 1.5E-02 |

**Table 37:** Adjusted P-values for pairwise T-tests for Quadratic Loss classifier evaluation on the Primary Fibroblasts IIC.\IY 120hpi dataset

|  | SVM | KNN | MAP |
| --- | --- | --- | --- |
| KNN | 1.7E-25 |  |  |
| MAP | 9.3E-32 | 1.9E-56 |  |
| MCMC | 9.3E-32 | 1.2E-54 | 7.1E-01 |

**Table 38:** Adjusted P-values for pairwise T-tests for Quadratic Loss classifier evaluation on the Primary Fibroblasts HCMV 96hpi dataset

|  | SVM | KNN | MAP |
| --- | --- | --- | --- |
| KNN | 6.5E-25 |  |  |
| MAP | 5.3E-32 | 1.1E-71 |  |
| MCMC | 7.1E-32 | 8.4E-71 | 5.7E-02 |

**Table 39:** Adjusted P-values for pairwise T-tests for Quadratic Loss classifier evaluation on the E14TG2a dataset

|  | SVM | KNN | MAP |
| --- | --- | --- | --- |
| KNN | 4.7E-04 |  |  |
| MAP | 4.7E-21 | 1.5E-103 |  |
| MCMC | 3.3E-12 | 1.8E-57 | 1.3E-137 |

### 5.7 Appendix 7: GO enrichment analysis figures

**Figure 16:**
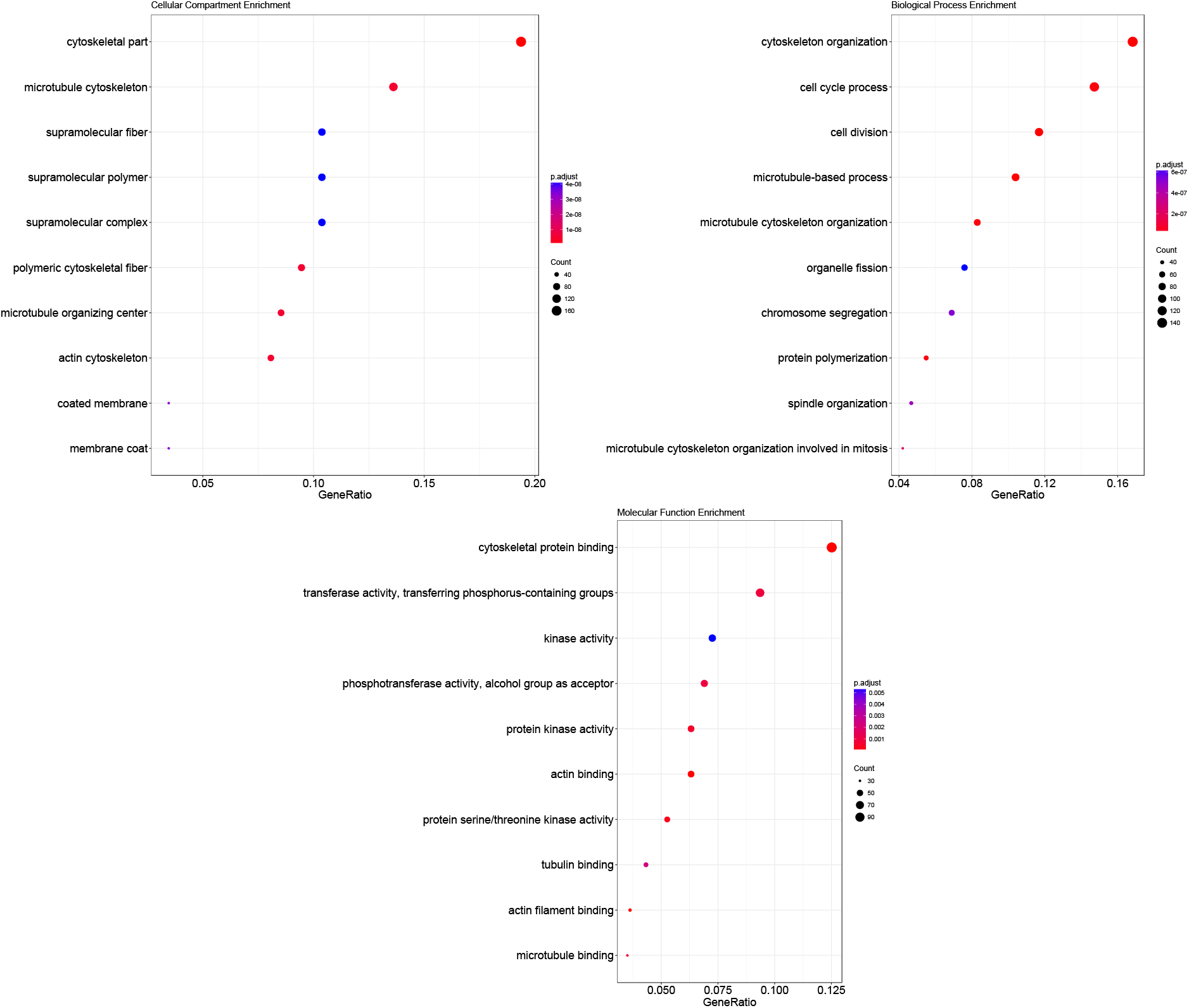
Gene Ontology over representation analysis on outlier proteins - that is proteins allocated with less than probability 0.95. We analyse the enrichment of terms in the cellular compartment, biological process, and molecular function ontologies. We display the top 10 significant results in the dotplots.

### 5.8 Appendix 8: Comparison of MCMC and MAP allocations

**Figure 17:**
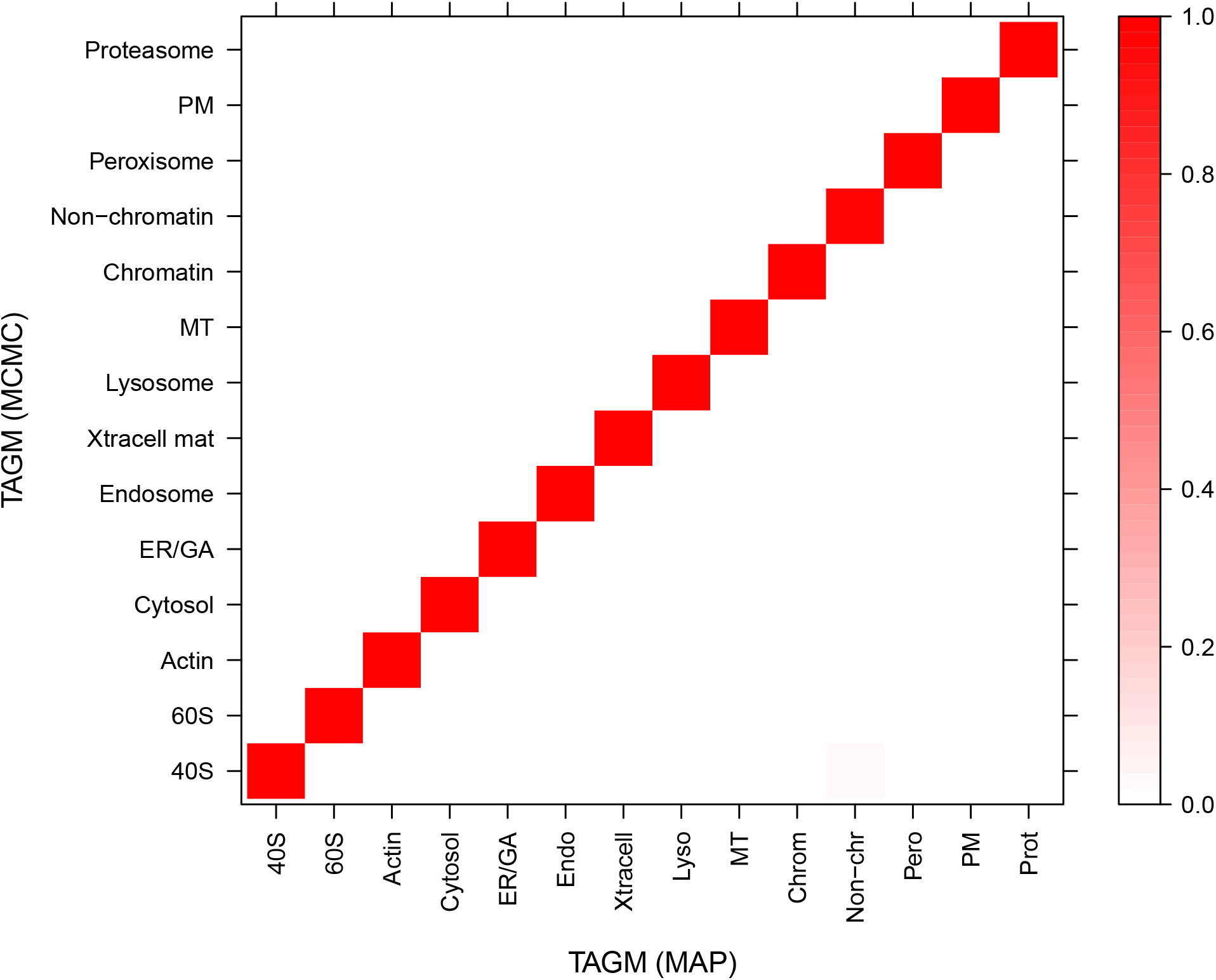
A heatmap representation of a contingency table comparing allocation produced by MCMC and MAP methods with posterior probability threshold set at 0.99 for both methods. The scale ranges from 0 to 1 with values indicating the proportion of assigned proteins to that sub-cellular location. Values along the diagonal represent agreement between classifiers whilst other values represent disagreement. The allocations of proteins by both methods are in strong agreement.

